# scGRIP: a graph-based explainable AI framework for single-cell multi-omics Gene Regulatory Inference with Prior Knowledge

**DOI:** 10.1101/2025.01.24.634773

**Authors:** Wenqi Dong, Manqi Zhou, Fei Wang, Yue Li

## Abstract

Multi-omics sequencing technologies can jointly measure transcriptome and chromatin accessibility at the single-cell resolution. This enables inference of gene regulatory networks (GRNs) at the cellular level, thereby elucidating high-resolution differential GRNs associated with diseases. However, existing methods lack interpretability and scalability. We present single-cell Gene Regulatory Inference with Prior knowledge (scGRIP), a graph variational autoencoder with 3 technical contributions. First, we treat transcription factors (TF), target genes (TG), and regulatory elements (RE) as nodes and their potential TF-RE and RE-TG interactions as edges using a prior cis-regulatory knowledge graph. Second, we tokenize the single-cell chromatin accessibility and gene expression with a shared codebook to compute cell-specific feature embedding. Third, we incorporate a GraphSHAP technique to infer GRN edge attribution at the single-cell level. scGRIP, evaluated against existing regulatory inference frameworks such as LINGER and scGLUE, demonstrates that integrating prior regulatory knowledge yields regulatory interaction features that improve discrimination of cell types and disease conditions. In particular, scGRIP identified coordinated activation of immune signaling, amyloid-related responses, and downstream target gene programs. The inferred GRNs for Alzheimer’s disease were supported by functional enrichment analyses and independent reference dataset.

## Background

Gene regulatory networks (GRNs) govern cellular identity and function by coordinating interactions among transcription factors (TFs), regulatory elements (REs), and target genes (TGs) to control gene expression programs.[1]. Understanding these regulatory relationships at single-cell resolution is beneficial for deciphering cellular heterogeneity, developmental trajectories, and disease mechanisms [2, 3]. The advent of single-cell sequencing technologies, such as single-cell RNA sequencing (scRNA-seq) paired with single-cell ATAC sequencing (scATAC-seq), has enabled the reconstruction of cell-specific GRNs by simultaneously profiling gene expression and chromatin accessibility within individual cells [4]. However, inferring accurate *cell-specific* GRNs remains challenging due to the sparsity, noise, and high dimensionality of the single-cell multiomic sequencing data [5].

Current multimodal GRN inference methods typically integrate multiple sources of regulatory information by predicting target gene expression from TF expression levels, associating TFs with accessible CREs via binding motif information [6], and linking CREs to target genes based on genomic proximity [7]. While methods like LINGER [6] have highlighted the value of incorporating external knowledge, the reliance on heterogeneous motif databases and diverse computational algorithms often leads to considerable variability across different frameworks. In parallel, advances in model interpretability have introduced principled attribution methods such as SHAP [8], which provide explanations for model predictions with strong axiomatic guarantees. Extensions of SHAP to graph neural networks, including GraphSVX [9], along with related approaches such as GNNExplainer [10], enable attribution of predictions to specific nodes and edges in graph-structured models. Recent approaches like scTFbridge [11] and LINGER [6] have begun to employ Shapley-value-based attribution to prioritize regulatory interactions. However, these techniques are often applied as post hoc tools rather than being fully integrated into GNN frameworks.

In this study, we present single-cell <u>G</u>ene <u>R</u>egulatory <u>I</u>nference with <u>P</u>rior knowledge (scGRIP), a framework that integrates graph neural networks (GNNs) [12], tokenization driven by the foundation-model [13], and graph-based Shapley value attribution [9] to infer cell-specific GRNs. First, scGRIP formalizes the regulatory landscape as a heterogeneous graph, treating TFs, REs, and TGs as nodes connected by potential interactions derived from cis-regulatory prior knowledge [7, 14]. Second, to transform this static topological scaffold into a dynamic model, we tokenize chromatin accessibility and gene expression profiles through a codebook [13] to compute cell-specific node embeddings. This allows scGRIP to capture the unique molecular state of each cell within the graph structure. Third, we leverage GraphSHAP [9] to perform Shapley-value-based attribution at the individual cell level. Consequently, scGRIP maps global biological priors onto individualized regulatory signatures, revealing heterogeneity and fine-grained dynamics that are typically masked by traditional integration methods.

Through comprehensive benchmarking on three multimodal single-cell datasets [15], we demonstrate that scGRIP outperforms state-of-the-art methods [6, 16] in cell-type-specific, cell-specific, and condition-specific GRN reconstruction tasks. The framework’s interpretable graph architecture and neighborhood aggregation strategy ensure both biological fidelity and computational scalability for large-scale datasets. Furthermore, the robustness of scGRIP’s learned representations is validated through downstream tasks, including cell-type clustering and cross-modality imputation. To showcase scGRIP in real-world applications, we conduct a case study on inferring differential cell-type-specific GRNs for Alzheimer’s Disease (AD). In particular, this enables the identification of disease-associated regulatory mechanisms, such as microglia-specific regulatory shifts in AD such as *APOE* and *SPI1*. Together, by enabling the direct quantification of TF-RE and RE-TG interactions in individual cells, scGRIP provides a framework for uncovering cell-type- and condition-specific regulatory programs.

## Results

### scGRIP model overview

scGRIP integrates single-cell multimodal data measured by paired scRNA-seq and scATAC-seq in the same cell using interpretable latent embeddings and external GRN databases (**Fig**. 1). Building upon the variational autoencoder [17] and multiomic Embedded Topic Model [18] (moETM), scGRIP combines multimodal data integration with gene regulatory network to incorporate prior biological information (**Fig**. 1a). Our contributions are threefold: graph-based representation learning, explainable AI for GRN inference, and model interpretability via topic modeling. First, we use node2vec [19] to generate static feature embeddings that reflect gene regulatory connectivity within the network, providing a structural view based on random walk-derived patterns (**Fig**.1c). Moreover, we adapted the xTrimoGene [13, 20] approach to learn cell-specific dynamic embeddings. To this end, we implement the efficient neighborhood aggregation strategy from GraphSAGE [21]. The final gene embeddings combine both node2vec and cell-specific embeddings, enriching node features with both immediate data-driven insights and broader network context. Second, we adapt GraphSVX [22], a graph SHAP explanation method for graph neural networks that extends SHAP from tabular data to graph data by treating neighboring nodes as contributors (i.e., TF and RE) to a target node (i.e., TG). Briefly, scGRIP repeatedly samples TFs or REs from a local k-hop GRN subgraph to construct a perturbed GRN by masking the values for TFs or REs and disconnecting their connections to the TG. It then fits a weighted linear surrogate model based on the Shapley kernel distance between the original and perturbed GRN so that the learned regression coefficients approximate the Shapley values, which can be interpreted as the contribution of each TF and RE to the TG (i.e., TF-TG and RE-TG links; **Fig**.1d). For TF-RE link, we take the dot product of their embeddings and average them across the cells of the same cell type to obtain the cell-type-specific TF–RE binding potential (**Fig**. 1 e). Third, we employ the ETM framework [23], which uses a linear decoder to enhance interpretability (**Fig**.1b). Building on our previous ETM-based methods [18, 24], the encoder enables the decoder to achieve accurate reconstruction by creating a linearly separable space via end-to-end training. The resulting linear decoder is parameterized only by the topic embedding, which is learned to align with the graph feature embedding.

**Figure 1:**
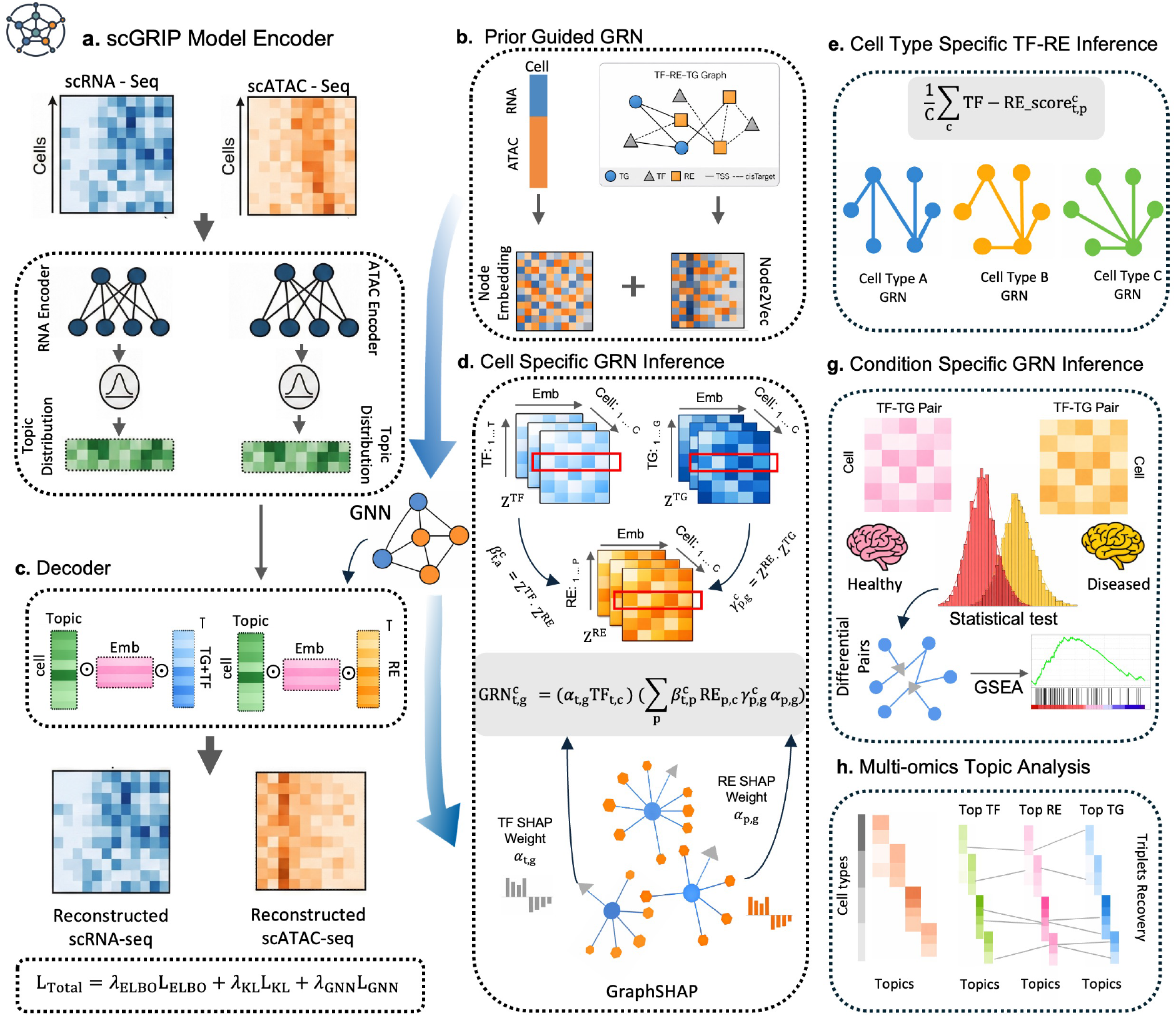
Overview of scGRIP. a. Multi-omics encoding. Single-cell RNA-seq and ATAC-seq count matrices are independently processed using modality-specific variational encoders to obtain low-dimensional cell-level topic distributions. **b. Prior-guided regulatory graph**. Prior biological knowledge is incorporated by constructing a TF–RE–TG regulatory graph based on transcription factor binding (cisTarget) and genomic proximity to transcription start sites (TSS). Node embeddings for TFs, REs, and TGs are learned via graph representation learning and integrated with the model to guide reconstruction. c. Graph-informed topic decoder. A linear embedded topic-model decoder integrates cell-level topic distributions with graph-derived embeddings to reconstruct both gene expression and chromatin accessibility, enabling joint modeling of TF, RE, and TG relationships across modalities. d. Cell-specific GRN inference. Regulatory interaction scores 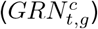 are computed at single-cell resolution by combining cell-specific TF and RE embeddings with node-level importance weights estimated using GraphSHAP, yielding interpretable regulatory contributions for each cell. e. Cell-type-specific GRNs. Cell-specific regulatory scores are aggregated across cells within each cell type, and statistical testing is applied to identify significant TF–RE interactions that characterize cell-type-specific regulatory networks. f. Disease-specific GRNs. Cell-specific regulatory interactions are stratified by condition and differential TF–TG relationships are identified through statistical testing. Significant regulatory differences are further characterized by downstream pathway enrichment analysis. g. Multi-omics topic analysis. Learned topics capture coherent biological programs across modalities. Cross-modality feature alignment enables recovery of TF–RE–TG triplets that reflect coordinated regulatory activity.

Building on these components, scGRIP enables three complementary downstream analyses of gene regulatory networks. First, cell-type-specific TF–RE inference (**Fig**. 1e) aggregates cell-level interaction scores across cells within the same cell type to estimate TF–RE binding potential, capturing motif-supported regulatory interactions that reflect cell-type-specific transcription factor binding preferences. Second, disease-specific GRN analysis (**Fig**. 1f) stratifies cell-specific regulatory interactions by condition and identifies differential TF–TG relationships through statistical testing, followed by pathway enrichment analysis to reveal condition-dependent regulatory mechanisms. Third, multi-omics topic analysis (**Fig**. 1g) leverages the interpretable topic structure to identify coherent regulatory modules, where top TFs, REs, and TGs within each topic are linked to form TF–RE–TG triplets, capturing coordinated regulatory programs across modalities.

### scGRIP accurately and efficiently infers cell-type-specific TF-RE relationships

We applied scGRIP to a single-cell PBMC multiome dataset and evaluated the cell-type-specific TF-RE prediction in terms of area under the precision-recall (AUPRC) to account for the class imbalance inherent in TF-binding site distributions. For the gold standard, we utilized the ChIP-Atlas database [25], which provides curated TF-binding peaks across 3,368 specific cell types or tissues. We compared scGRIP against two state-of-the-art methods, LINGER [6] and GLUE [16], as well as a baseline Pearson correlation coefficient (PCC) between gene expression and chromatin accessibility across cells at the cell-type level. scGRIP consistently demonstrated better performance, achieving an average AUPRC of 0.613 across 14 TFs and 3 cell types, outperforming GLUE (0.606), LINGER (0.599), and the PCC baseline (0.588) (**Fig**. 2a, **Table** S1). Specifically, for CD4 T cells on average, scGRIP reached 0.630, while LINGER scored 0.620 and GLUE achieved 0.625; in B cells, scGRIP outperformed both with a score of 0.610, compared to LINGER (0.585) and GLUE (0.605); in memory B cells, scGRIP scored the highest score of 0.620, while LINGER averaged 0.615 and GLUE achieved 0.605. These results underscore the ability of scGRIP to consistently outperform other methods in inferring cell-type-specific TF binding sites. We then focused on the predicted regulatory landscapes for a specific TF, BCL6. scGRIP predicted binding sites in B cells with an AUPRC of 0.650, substantially higher than LINGER (0.580). Indeed, we observe higher prediction scores within experimentally validated TF-binding regions (**Fig**. 2b, red box). We also calculated PCC for the top 3000 TF–RE pairs ranked by LINGER, of which 137 TF–RE pairs with significant PCC with average PCC equal to 0.29. In contrast, scGRIP identified 217 BCL6–RE pairs with significant PCC with average PCC equal to 0.32. Among the evaluated pairs, 104 of the 137 LINGER predictions overlapped with ChIP-seq peaks, compared with 217 of the 149 scGRIP pairs. Beyond predictive accuracy, scGRIP offers architectural and computational advantages. Specifically, scGRIP utilizes a GraphSAGE-based architecture to scale with graph size (**Fig**. S1). This allows scGRIP to learn the entire GRN from single-cell multiome data in a unified framework while maintaining high memory efficiency. In contrast, LINGER trains one model per target gene and 20K genes for all 20K genes; GLUE needs to subsample nodes as the graph grows to keep memory in check, but this comes at the cost of performance (**Fig**. S1).Overall, scGRIP improves cell-type-specific TF–RE inference by recovering more ChIP-seq-supported regulatory pairs with higher precision and stronger concordance with chromatin accessibility patterns.

**Figure 2:**
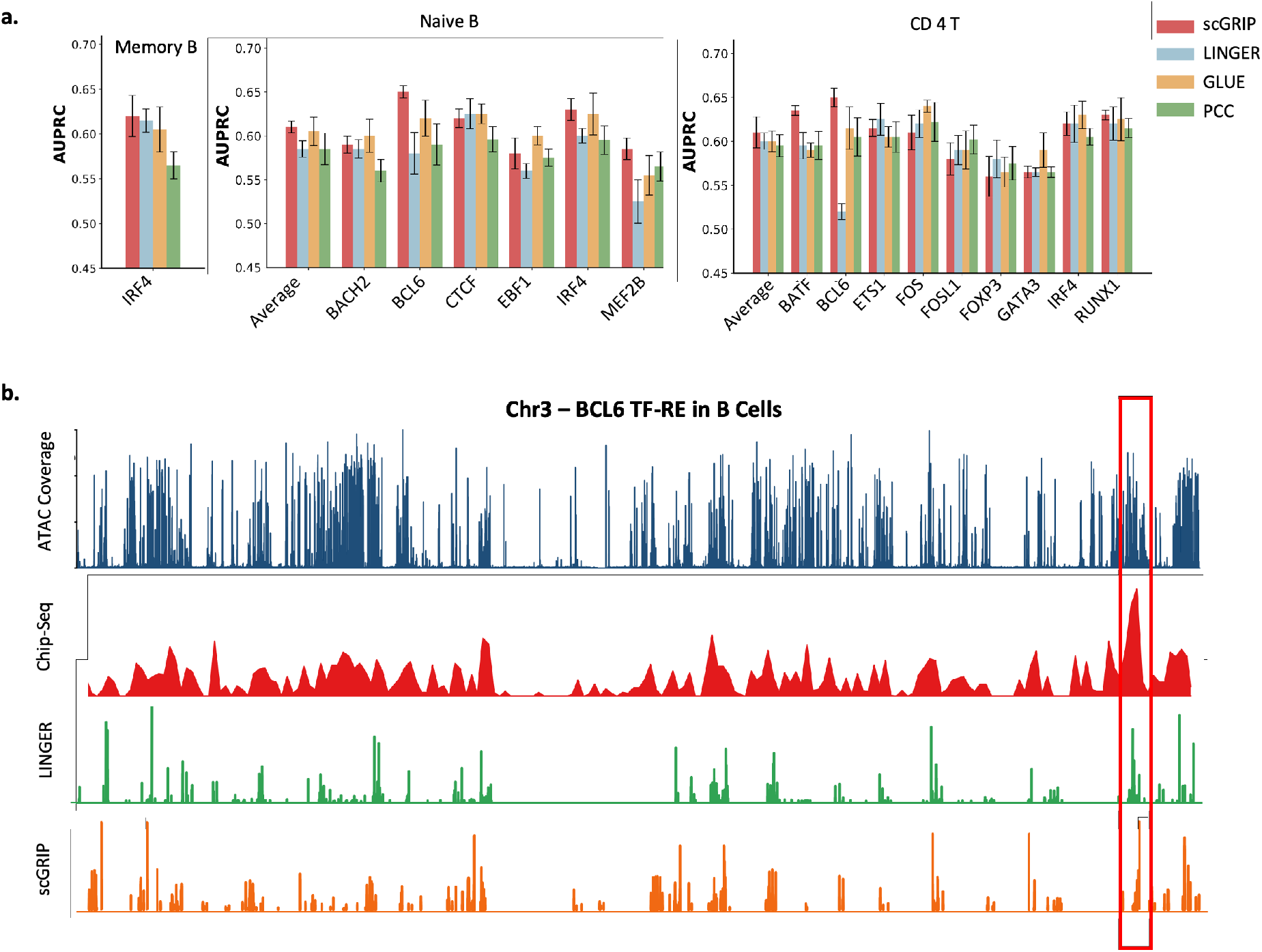
Evaluation of cell-type-specific TF-RE inference using PBMC multiomes. **a.** Comparison of TF-RE binding potential inference across cell types and methods. Bar plots show the AUPRC for scGRIP, LINGER, GLUE, and PCC, evaluated across TFs, with error bars indicating variability across independent training runs. **b**. Locus-level comparison of TF-RE binding predictions for BCL6 in B cells on chromosome 3. From top to bottom: chromatin accessibility signal in B cells measured by scATAC-seq; BCL6 ChIP-seq binding sites from ChIP-seq Atlas; TF-RE binding potential predicted by LINGER; and TF-RE binding potential predicted by scGRIP. The highlighted region indicates concordant chromatin accessibility, ChIP-seq support, and comparable scGRIP prediction to LINGER.

### scGRIP-inferred GRNs accurately predict cell types

We evaluated scGRIP-inferred cell-specific regulatory potentials in capturing distinct cell types by performing binary cell-type classification on an Alzheimer’s disease (AD) multiome dataset [15]. To avoid additional model training, we used the instance-based classifier *k*-nearest neighbor (KNN), which uses the predicted regulatory interaction scores between TF and TG as the input features to predict cell types. KNN on scGRIP-inferred GRNs achieved consistently strong AUROC across cell types (**Fig**. 3a). We compared scGRIP against LINGER and a SHAP-ablation variant of scGRIP, in which regulatory potentials are computed directly from the dot product of learned embeddings without SHAP-based attribution weighting. Across individual cell types scGRIP demonstrated consistently stronger TF–TG recovery compared with both the ablation variant lacking SHAP-derived attribution weights and the prior-constrained framework *LINGER*. In astrocytes and excitatory neurons, scGRIP achieved the highest AUROC. A similar trend was observed in inhibitory neurons and microglia, where the performance gap over the ablation model and the more moderate margin over *LINGER* indicate that SHAP-informed edge weighting helps capture subtle, context-dependent regulatory interactions while refining core immune and neuronal regulatory signals. For OPCs and oligodendrocytes, scGRIP maintained stable improvements or comparable performance relative to *LINGER*. The results indicate that incorporating SHAP-derived attribution weights is an important step for capturing cell-specific regulatory dynamics.

**Figure 3:**
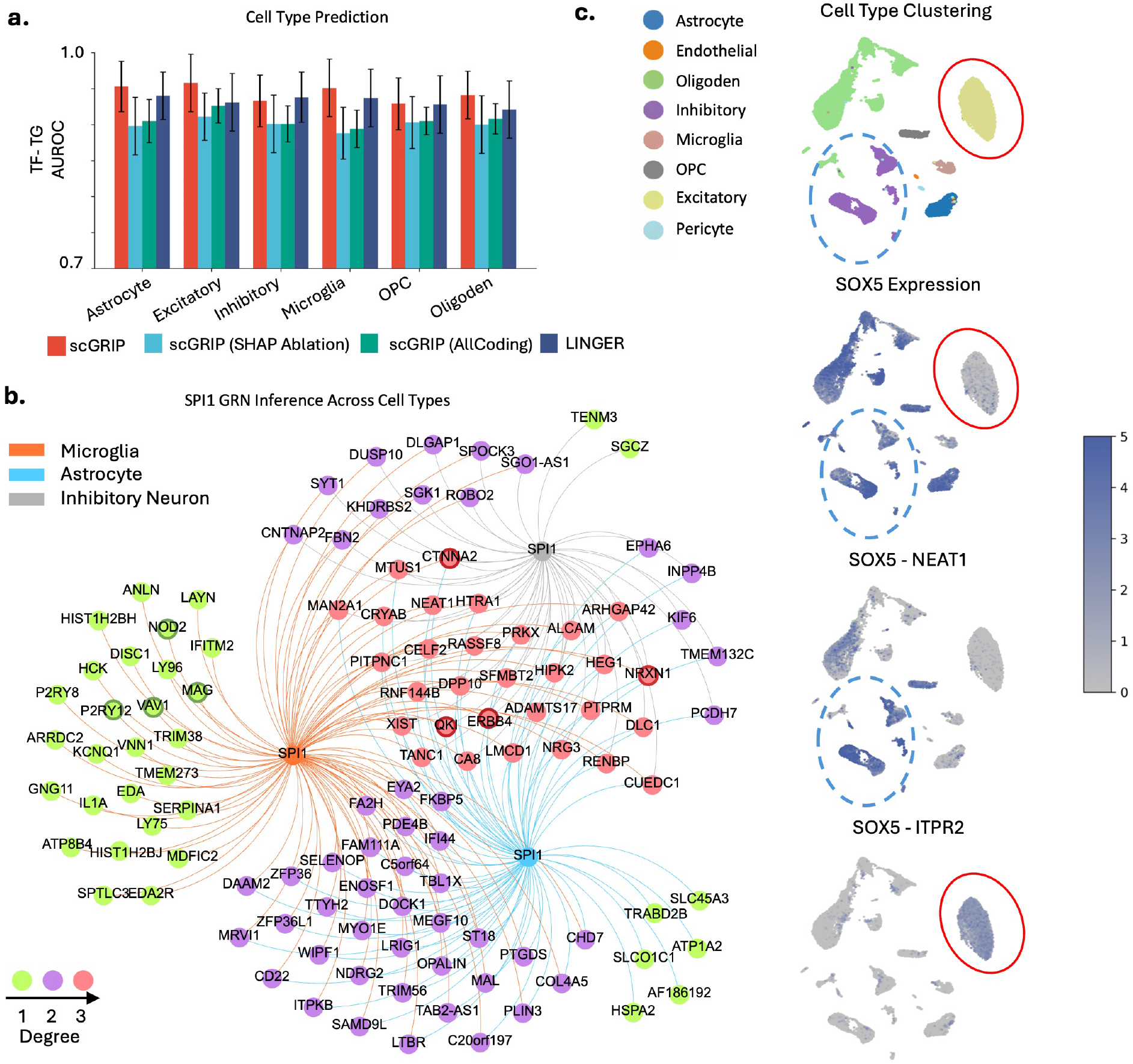
scGRIP infers cell-level regulatory activity. **a.** Cell-level classification performance for predicting cell types using KNN based on elements of TF–RE interaction scores. Bar plots show the AUROC across cell types, with error bars indicating variability across cross-validation runs. **b**. Cell-specific regulatory networks of SPI1 across cell types. Node color indicates node degree, reflecting regulatory connectivity. Key regulatory nodes are highlighted with grey outlines. Edge colors denote the specific cell type where the interaction is active. **c**. UMAP of cellular state and regulatory activity. Cells are projected into a shared latent space via UMAP based on their learned cell embedding. Overlaid color intensity represents either *SOX5* expression or specific cell-level regulatory activity scores (*SOX5*(TF)–*NEAT1*(TG) and *SOX5*(TF)–*ITPR2*(TG)).

We then examined the aggregated cell-type-specific networks derived from the underlying cell-specific scGRIP-inferred GRNs. We focused on SPI1 across multiple cell types as it is a lineage-defining TF of the ETS-family that plays a central role in hematopoietic and myeloid differentiation, including microglial development and the regulation of immune gene in the brain [26]. In microglial cells, we discovered several microglia-specific SPI1 target genes (*NOD2, MAG, VAV2, P2RY12*) (the dark green–outlined nodes in **Fig**. 3b). These canonical innate immune regulators are primarily involved in immune receptor signaling and phagocytosis, providing biological support for the inferred regulatory structure [27]. Furthermore, signaling mediators such as *HCK* are characteristic of myeloid lineages [28–30]. The recovery of these targets indicates that scGRIP highlights biologically meaningful regulatory interactions at the single-cell level. Many additional targets predicted to be regulated across the three cell types are genes involved in synaptic signaling and neuronal connectivity (red nodes in **Fig**. 3b). These include components of neuregulin signaling (*ERBB4* [31, 32]) and synaptic adhesion (*CTNNA2* [33]). Their presence across multiple cell types likely reflects shared regulatory programs affecting synaptic maintenance and neuronal communication, rather than strict cell-type specificity. Such patterns are consistent with neurodegenerative disease contexts, where alterations in synaptic function and neuronal remodeling emerge as broadly coordinated responses across diverse cellular populations [34].

We then determined whether these inferred potentials reflect TF activities or capture independent regulatory logic by comparing *SOX5* expression levels with cell-specific regulatory strengths across cell types. We observe that *SOX5* expression is highly specific to inhibitory neurons and nearly absent in oligodendrocytes (**Fig**. 3c). Its scGRIP-inferred regulatory activity follows a more nuanced pattern: *SOX5* (TF)–*NEAT1* (TG) regulatory interaction is highly active in inhibitory neurons where *SOX5* is abundantly expressed, but inactive in oligodendrocytes. Conversely, the *SOX5*(TF)–*ITPR2*(TG) interaction exhibits strong regulatory activity specifically in oligodendrocytes despite minimal *SOX5* transcript levels, while remaining silent in inhibitory neurons. Thus, scGRIP leverages dynamic node features to infer cell-specific regulatory interactions that capture functional cell states. The inferred links encode regulatory programs sufficient to distinguish cell types, indicating that cellular identity is reflected not only in transcript abundance but also in context-dependent regulatory activity.

### scGRIP reveals condition-specific GRNs in Alzheimer’s disease

We applied scGRIP to dissect AD-associated regulatory potentials within specific cellular contexts (**Fig**. 1 g). To this end, we performed differential GRN inference between AD and control samples at the single-nucleus resolution using the same data as benchmarked above [15]. We focused our analysis on microglia, a key cellular mediator of neuroinflammation [35]. scGRIP revealed significant regulatory relationships in AD microglia, identifying condition-specific TF–TG interactions using a two-sided Wilcoxon rank-sum test with Benjamini–Hochberg FDR correction, with significance defined at an adjusted *P <* 0.05. Among the most significantly altered pairs, 37% of target genes overlapped with DEGs, capturing well-characterized AD-associated genes and microglial activation markers (**Fig**. 4a). For instance, *APOE*, a primary genetic risk factor for late-onset AD [36], emerged both as an up-regulated target gene in the TF–TG pair *TREM2*–*APOE*. Similarly, *CD86*, a marker of microglial activation [37], was up-regulated at both the expression and regulatory levels through the *HCLS1*–*CD86* interaction. Conversely, *CXCL10*, a gene coding for chemokine protein involved in microglial homeostasis, appeared as a down-regulated target gene by the TF *IRF8*. These top-ranking pairs exhibited a bifurcation between disease states, where AD samples displayed distinct regulatory activity patterns compared to controls. Specifically, pairs such as *ZNF98*–*APOE* showed consistently higher regulatory potentials in AD (**Fig**. 4b), reflecting a coordinated network-level activation.

**Figure 4:**
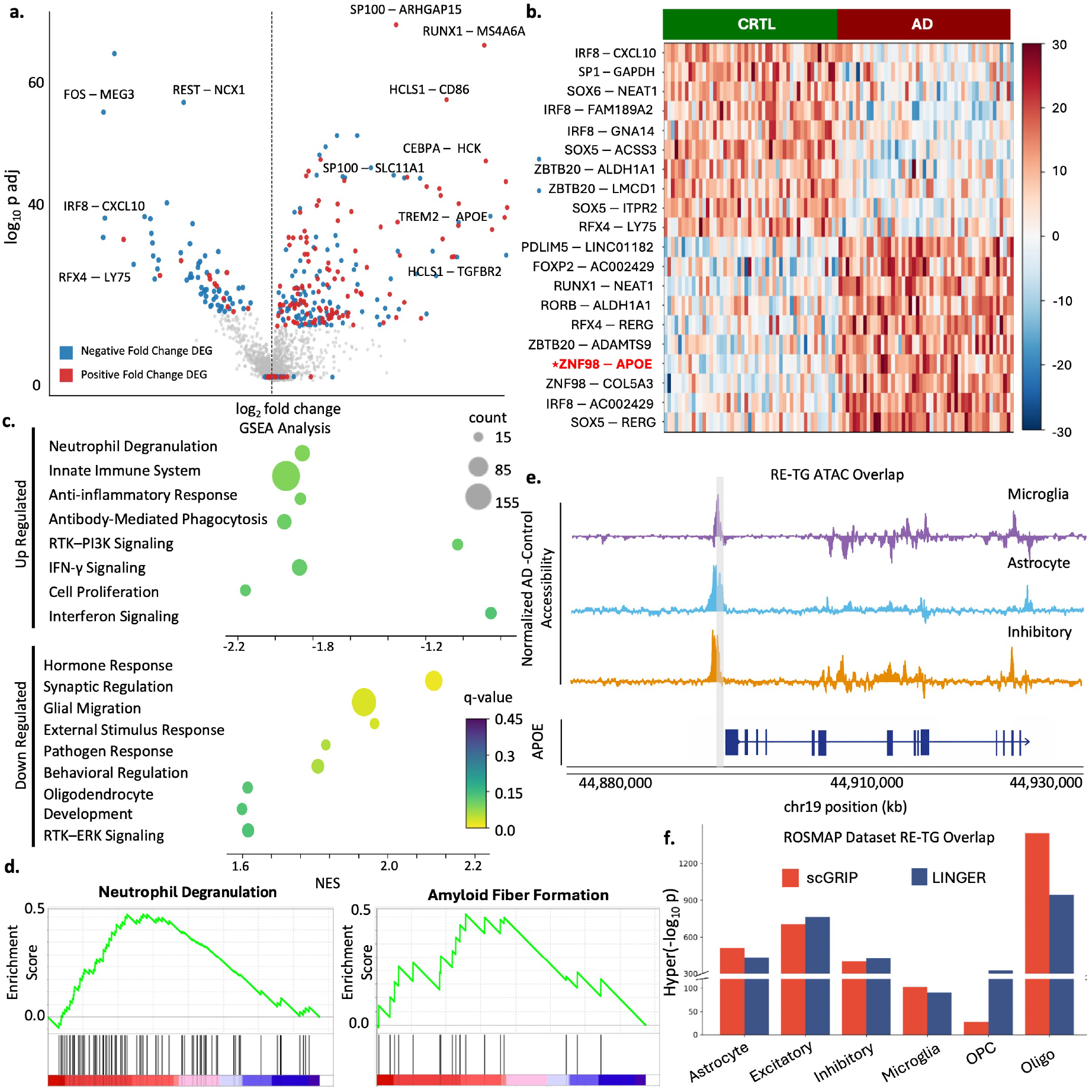
scGRIP infers condition-specific regulatory programs in Alzheimer’s disease. **a.** Condition-specific differential regulatory activity in microglia. The volcano plot displays significant condition-specific TF–TG interactions, with red and blue indicating overlaps with positively and negatively fold-changed DEGs, respectively. **b**. Condition-specific regulatory shifts. The heatmap illustrates the top-ranking upregulated TF–TG pairs in AD microglia compared to controls, derived from inferred regulatory interaction scores. **c**. Gene set enrichment analysis of target genes from significant TF–TG pairs identified in microglia. **d**. Representative GSEA enrichment plots for selected pathways associated with AD regulatory programs. **e**. Genomic locus example illustrating RE–TG inference for *APOE*. The top three tracks show AD–control chromatin accessibility differences in microglial cells, astrocytes and inhibitory neurons, and the bottom track indicates the genomic location of *APOE*. The predicted regulatory element (grey bar) lies within the TSS region of *APOE* and overlaps with a major accessibility peak. **f**. Comparative validation of regulatory links. Hypergeometric enrichment scores quantifying the overlap of inferred RE–TG pairs with the ROSMAP validation dataset results, comparing scGRIP with the baseline method LINGER.

To characterize the functional landscape of these disease-associated regulatory networks, we performed Gene Set Enrichment Analysis (GSEA) [38] on the target genes of the significant TF–TG pairs. The up-regulated transcriptional programs in AD microglia were significantly enriched in pathways such as neutrophil degranulation, innate immune activation, adjusted p-value 0.017, and interferon-gamma signaling (**Fig**. 4c). Specifically, the enrichment of neutrophil degranulation is closely associated with the neuroinflammatory environment of the AD brain [39]. Furthermore, the activation of amyloid-related pathways, as evidenced by the GSEA leading-edge curve for amyloid fiber formation (**Fig**. 4d), reflects a specialized microglial response to amyloid-*β* accumulation [40, 41]. These findings align with the established AD disease models, where the accumulation of amyloid fibers and the excessive activation of neutrophil degranulation create a toxic pro-inflammatory environment [42, 43]. This environment actively suppresses the synaptic gene regulation, with synaptic loss serving as a primary predictor of cognitive decline [44, 45]. Our observation of up-regulated neutrophil degranulation and amyloid fiber formation, paired with the down-regulation of synaptic regulation, is highly consistent with these mechanisms. These results suggest that the condition-specific regulatory potentials inferred by scGRIP accurately reflect established disease-driving biological processes. To provide mechanistic evidence for these condition-specific predictions, we examined the epigenomic landscape of the *APOE* locus. scGRIP predicted a high-confidence cis-regulatory element located within the TSS region of *APOE* that coincided with a AD-specific chromatin accessibility peak (**Fig**. 4e). Notably, this RE–TG interaction was consistently recovered across these distinct cell types, demonstrating the model’s ability to identify robust regulatory links that align with physical epigenetic shifts.

Finally, we benchmarked scGRIP predictions against another AD dataset [46], which provides an independent resource of single-nucleus chromatin accessibility and gene expression from postmortem human brain tissue. Based on hypergeometric enrichment analysis, we quantified the overlap between scGRIP-inferred RE–TG pairs and ROSMAP-validated regulatory interactions across six major brain cell types (**Fig**. 4f). scGRIP achieved higher enrichment than the LINGER baseline in astrocytes, microglia, and oligodendrocytes, with comparable performance in excitatory and inhibitory neurons (**Fig**. 4f). Although LINGER showed stronger enrichment in OPCs, scGRIP predictions remained statistically significant across all six cell types. Collectively, these findings demonstrate that scGRIP accurately captures the complex, condition-specific regulatory architecture underlying AD, offering validated predictions that bridge transcriptional networks and the AD etiology.

### scGRIP-inferred topics capture GRN submodules

Given the interpretable topic-model decoder, scGRIP can automatically identify coherent regulatory modules within the learned topic distributions at the both cell level and feature level. For example, we identified topics related to microglia cells (21, 24, 44, 92, and 61). At the cell level, the corresponding topic mixtures exhibit clear enrichment within microglia populations and more distinct patterns across micrgrolia subtypes compared to the graph-ablated variant (**Fig**.5a). At the feature level, the top TFs, REs, and TGs within each of these topics exhibit coordinated regulatory patterns with high between-modality TF-RE and RE-TG connectivity within the same topic (**Fig**.5b). These results suggest that incorporating prior regulatory graph information can resolve finer-grained cellular heterogeneity and capture biologically structured regulatory signals. We found TFs associated with immune and myeloid lineage regulation among the top TFs within the 5 topics.

**Figure 5:**
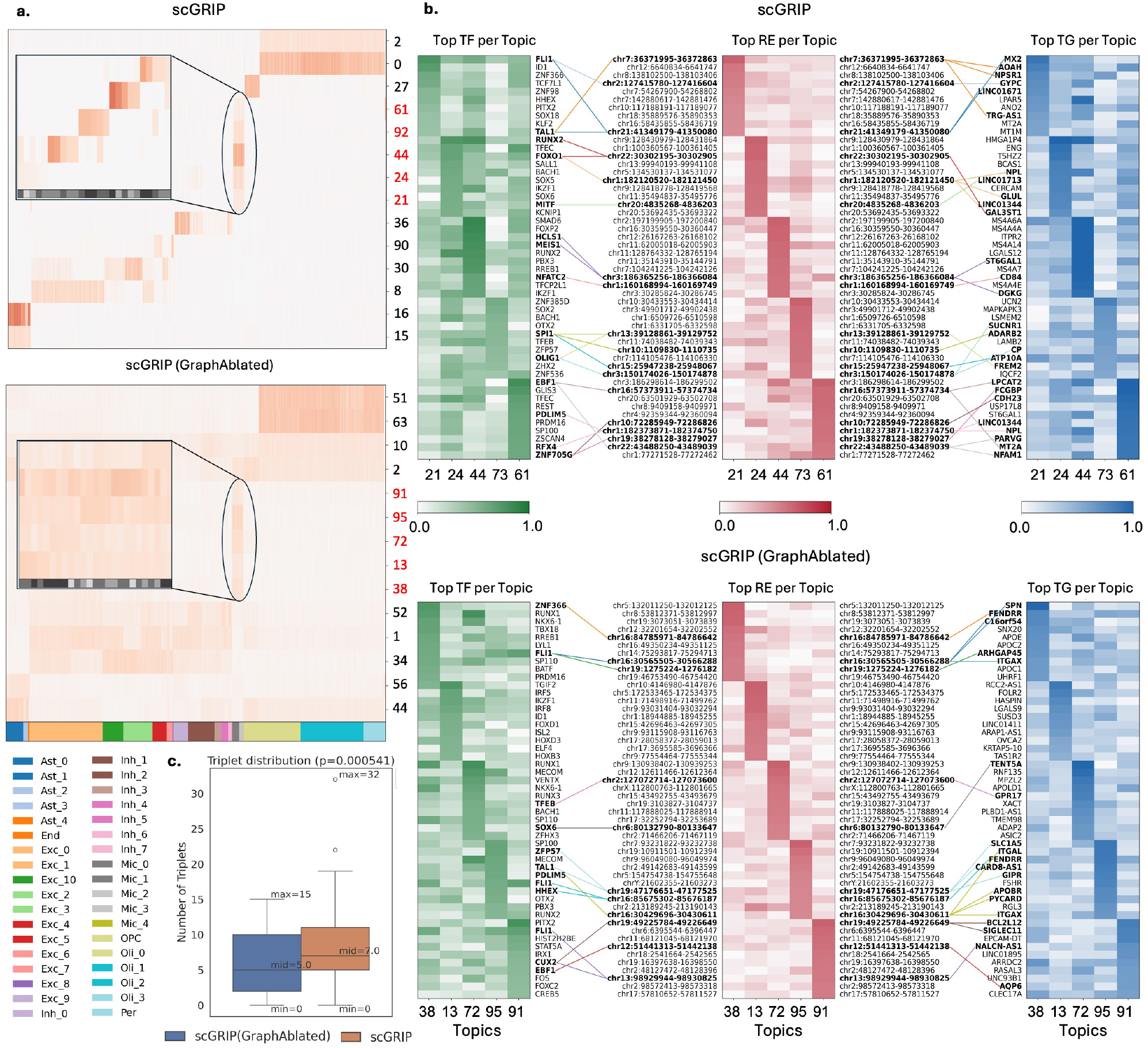
Topic analysis reveals enriched regulatory modules in the AD dataset. **a.** Heatmaps showing the intensity of the top topics from the AD dataset using scGRIP (top) and scGRIP (Graph-ablated) (bottom) across cell types. Topics highlighted in red correspond to microglia-associated topics. The inset sub-heatmaps show the top five cells per topic within microglia subcell types, illustrating the distinct topic weight patterns within microglial populations. **b**. Feature analysis for the selected topics. Heatmaps display the top 10 transcription factors (TFs), regulatory elements (REs), and target genes (TGs) ranked by topic weight. TF–RE–TG regulatory triplets identified within each topic are highlighted with colored connections, illustrating topic-specific regulatory modules captured by scGRIP. Bold labels denote TFs, REs, and TGs that participate in the identified regulatory triplets. **c**. Comparison of the number of regulatory triplets identified among the top features per topic between scGRIP and the graph-ablated variant. scGRIP identifies a greater number of regulatory triplets, indicating improved recovery of coherent regulatory modules when incorporating graph-based regulatory priors.

NFATC2 under topic 44 is a calcium-responsive TF of the NFAT family known to regulate immune activation and signaling pathways in both lymphoid and myeloid cells [47]. scGRIP identifies *CD84*, a SLAM family receptor involved in immune cell signaling, as a downstream target gene, further supporting the enrichment of key characteristics of activated microglia in neuroinflammatory conditions [48]. *SPI1* under topic 73 is a master regulator of myeloid cell identity and is essential for microglial development and function [49]. scGRIP identifies accessible RE bound by *SPI1* (for example, chr10:1109830-1110735) that connect to downstream target genes including *ADARB2*, a neuron-specific gene that modulates RNA editing [50]. The triplet reflects the coordinated activity across regulatory programs. Co-occurence of *FREM2*, a protein involved in cell matrix interaction, and *SPI1* reflects downstream extracellular matrix changes associated with neuroinflammation and microglial activation that enable immune functions such as phagocytosis [51] (**Fig**. 5b).

Furthermore, examining the relationships among the top features of each topic reveals numerous TF-RE-TG triplets. While some topics do not contain detectable TF–RE–TG triplets among their top features, scGRIP consistently identifies a greater number of regulatory triplets overall. In particular, both the median and the maximum number of recovered triplets per topic increase compared to the graph-ablated variant, indicating that incorporating graph-based regulatory priors improves the model’s ability to reconstruct coherent regulatory relationships (**Fig**.5c). In PBMC dataset, a similar result is observed (**Fig**.S4), where The graph ablated variant recovered significantly fewer numbers of the prior relationship than the proposed scGRIP.

To further investigate topic related TF-TG regulatory activity, we inferred directly the active TF-TG regulatory interactions by calculating average TF-TG regulatory scores across the top 10% of cells with the highest weights for each topic. TF-TG pairs with scores above the 95th percentile are considered active. TF-TG scoring revealed several putative regulatory associations. Specifically, TF *NFATC2* is connected to genes from the *MS4A* locus, including *MS4A6A* and *MS4A4E*, which are established microglia marker genes associated with AD [52] (**Fig**. S3), providing additional evidence for inflammatory regulatory program captured in topic 44, consistent with the patterns observed in the triplet analysis.

### scGRIP confers superior cell representations and cross-modality imputation

As a standard benchmark, we evaluated cell-level representation learned by the scGRIP encoder based on cell clustering metrics. We compared scGRIP against state-of-the-art multi-omics integration methods, such as moETM [18], MultiVI [53], and Seurat V4 [54], across three independent datasets. Overall, scGRIP matched the top-performing method across all four evaluation metrics (**Fig**. 6a, b, **Table** S2). UMAP visualizations further confirmed these results, as scGRIP maintained clear separation between neighboring populations, such as ‘B1 B’ and ‘CD 16+ Mono’cells, where competing methods showed significant overlap (**Fig**. 6c). Ablation studies revealed that excluding the GRN component (scGRIP-noGraph) consistently degraded performance, highlighting the importance of regulatory graph structures in learning discriminative cell-type identities. Furthermore, we evaluated the model’s ability to perform cross-modality translation, specifically predicting gene expression from chromatin accessibility (ATAC2RNA) and vice versa (RNA2ATAC). To optimize these tasks, we fine-tuned scGRIP, which incorporates a specialized BABEL-style imputation loss [55]. scGRIP outperformed existing models, including BABEL, achieving an average Pearson correlation of 0.73 for ATAC2RNA and 0.57 for the more challenging RNA2ATAC task (**Fig**. 6d-h, **Table** S3). Ablation experiments demonstrated the addition of the imputation loss provided consistent gains in accuracy across all gene sets (**Table** S3). Qualitatively, the imputed values maintained strong linear correlations with observed data, demonstrating that scGRIP accurately captures the functional interplay between epigenetic states and transcriptome.

**Figure 6:**
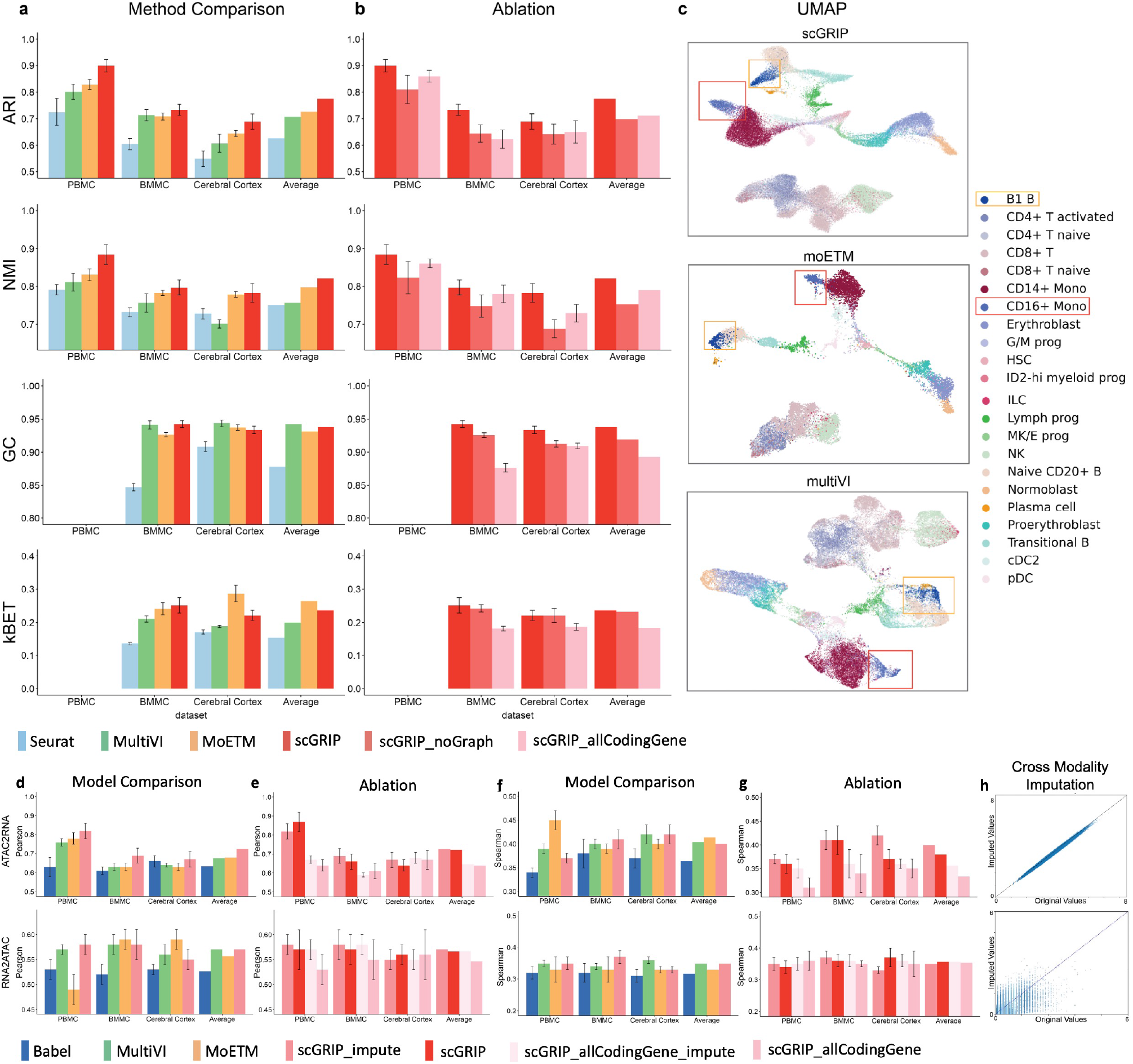
Methods comparison based on cell clustering and modality imputation. **a** Individual performance of each method on each dataset, as well as the averaged values across all datasets. Each row corresponds to a different evaluation metric. Since the PBMC dataset consists of only one batch, the batch effect removal evaluation metrics, GC and kBET, were not applicable and are therefore left blank for the PBMC dataset. **b** Performance of scGRIP with its ablated versions. **c** UMAP visualization on the BMMC dataset. **d-g** Methods comparison based on cross-modality imputation. The upper panel displays performance on the ATAC2RNA imputation task, while the lower panel shows performance on the RNA2ATAC imputation task. **d**,**e** Pearson correlation for each method and scGRIP’s ablated versions on each dataset, along with the average Pearson correlation values across all datasets. **f**,**g** Spearman correlation for each method and scGRIP’s ablated versions on each dataset, along with the average values across all datasets. **h** Scatterplot of original versus imputed values, with the diagonal line shown in blue.

## Discussion

Inferring cell-specific GRN becomes possible with the advent of single-cell multi-omic technologies. However, existing methods are limited in accurate GRN inference at the single-cell resolution. In this study, we develop scGRIP, a graph-based framework designed to infer cell-specific GRNs from single cell multi-omics data. By incorporating prior regulatory knowledge and dynamic cell embeddings, scGRIP focuses on uncovering cell-resolved regulatory networks. The improved GRN inference performance is enabled by three key contributions: 1) scGRIP constructs a cis-regulatory knowledge graph where TFs, REs, and TGs are represented as nodes connected by potential TF–RE and RE–TG interactions; 2) scGRIP tokenizes single-cell chromatin accessibility and gene expression data through a shared codebook to generate cell-specific node embeddings; 3) scGRIP employs GraphSHAP-based technique to quantify edge importance and infer GRNs structures at the single-cell level.

Analysis of cell-type-specific gene regulatory networks indicates that inferred TF–RE interactions are strongly supported by known transcription factor binding motifs. Using ChIP-seq Atlas as ground truth, scGRIP achieves high AUPRC in recovering TF binding sites across cell types, consistent with experimentally validated binding patterns. scGRIP leverages dynamic node features to infer cell-specific regulatory networks. By integrating gene expression and chromatin accessibility into these embeddings, the scGRIP captures context-dependent regulatory interactions at single-cell resolution. The inferred links encode cell-specific transcription factor activity and downstream target regulation that enable cell-type discrimination, demonstrating that regulatory interactions capture cellular identity beyond gene expression alone. Extending this framework to disease contexts, scGRIP identifies coordinated regulatory shifts in microglia in Alzheimer’s disease, including *SPI1* and *APOE* associated interactions that reflect activation of immune and amyloid-related pathways.

While the use of GraphSAGE for neighborhood sampling provides good scalability for large biological networks, our memory consumption analysis reveals competitive performance. scGRIP demonstrates reasonable memory scaling as graph size increases, performing significantly better than attention-based approaches such as GAT (**Fig**.S1). Moreover, our method achieves comparable memory scaling characteristics to GLUE, which also employs efficient subgraph training strategies, making scGRIP suitable for processing large-scale single-cell multi-modal datasets without excessive memory requirements [56].

Despite these advances, several limitations remain. First, the construction of regulatory edges in scGRIP relies on a fixed genomic proximity threshold to the TSS, without differentiating between upstream and downstream distances. While this proximity-based strategy is widely used and provides a scalable first approximation, previous studies suggest that regulatory potential may vary directionally, potentially affecting assignment accuracy [57]. Second, relying solely on linear genomic distance may fail to capture long-range enhancer–promoter interactions mediated by three-dimensional chromatin architecture, which are increasingly recognized as critical for gene regulation [58, 59]. Incorporating chromatin conformation data, such as Hi-C [60, 61], into future versions of scGRIP could further refine regulatory element–gene linkages by accounting for spatial proximity.

As future work, we plan to explore several directions. First, we will explore transformer-based graph convolution to uncover de novo regulatory interactions not captured in existing GRN graph databases. Second, other omics data types such as single-cell proteomics could further improve downstream interpretability. This can be done in a mosaic data integration. Third, we will extend scGRIP to model spatial transcriptomic data by identifying spatiotemporal-specific regulatory circuits and cell-cell communications via ligand-receptor interaction network [62].

## Conclusion

In summary, we present scGRIP, an interpretable and scalable framework for inferring cell-specific gene regulatory networks from single-cell multi-omics data. By integrating prior regulatory knowledge with dynamic embeddings and graph-based attribution, scGRIP captures regulatory interactions that define cellular identity and disease states consistent with state-of-the-art methods. In Alzheimer’s disease, scGRIP uncovers microglia-specific regulatory programs associated with immune activation and amyloid-related pathways.

## Methods

### Prior GRN construction

The prior GRN is a graph *G* = (*V, E*), where TFs, REs, and TGs are nodes *V* and their regulatory interactions as edges *E* between nodes. Each edge *e*_*ij*_ ∈ *E* reflects a regulatory interaction between two nodes *v*_*i*_, *v*_*j*_ ∈ *V*. Although REs are generally defined as functional regulatory regions such as enhancers or promoters [63], we consider each accessible peak identified from scATAC-seq data as an RE to ensure genome-wide coverage. For TF-RE, we add an edge *e*_*tp*_ between a TF *t* and a RE *p* if the cisTarget [7] database predicts that TF *t* binds motifs within the peak *p*. For RE-TG, we link an RE *p* to a TG *g* if the genomic distance between the transcription start site (TSS) of *g* and the start position of *p* is within a predefined threshold *d*_TSS_, set to 1 Mbp based on previous study [14]. The resulting undirected graph *G* is represented by an adjacency matrix *A* of dimension (*M* × *M*), where 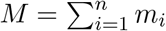 represents the total number of nodes, specifically the combined number of genes and peaks across modalities. In the current framework, TFs are treated as a subset of genes, and therefore are not explicitly represented as separate node types in the graph. Only interactions between different types of entities are considered, while the diagonal blocks corresponding to gene-gene and RE-RE interactions are excluded.

### Cell-specific dynamic node features for the GRN

For the initial node feature embedding, we first train a node2vec [19] model to capture the global structural properties of the GRN. Specifically, node2vec generates embeddings that reflect node connectivity within the network, providing a macroscopic view based on random-walk derived patterns. These pre-trained node2vec embeddings are integrated with the cell-specific embeddings via xTrimoGene [13, 20], which transforms each gene expression scalar value into a latent vector as follows:

1. Project the scalar value 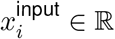 for gene *i* onto score vector **h**_*i*_ ∈ ℝ^*b*^:

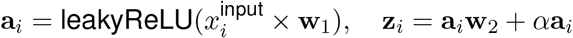

where **w**_1_ ∈ ℝ^1×*b*^, **w**_2_ ∈ ℝ^*b*×*b*^, and *α* are learnable parameters and **a**_*i*_ ∈ ℝ^*b*^, and **z**_*i*_ ∈ ℝ^*b*^ are b-dimensional vectors.
2. Normalize **z**_*i*_ with softmax to produce a vector **γ**_*i*_ ∈ ℝ^1×*b*^:

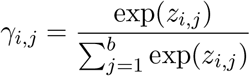
3. The value embedding is a weighted sum of all *b* embeddings from the learnable embedding codebook **T** ∈ ℝ^*b*×*d*^:

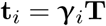

where **t**_*i*_ ∈ ℝ^1×*d*^. Zeros are encoded **t**_*i*_ = **0** if the gene *i* is not expressed in the cell.
4. The final input embedding for gene *i* is:

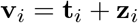

where 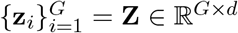 denotes node2vec embedding for all *G* genes.

In the context of token embedding in NLP, the gene-independent and value-dependent latent vector **t**_*i*_ can be considered as the ‘position embedding’ and the gene-specific and value-independent node2vec embeddings 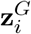 are the ‘word embeddings’. The process culminates with the final gene embeddings being a combination of attention-adjusted features and node2vec embeddings, thus enriching the node features with both data-driven insights and broader network context. Analogously, peak embeddings are obtained by applying the same value encoder to peak accessibility values and combining them with peak-specific node2vec structural embeddings, followed by GNN propagation to produce cell-specific peak representations.

### Graph neural network component

We employed a neighborhood aggregation strategy inherited from GraphSAGE [21]. GraphSAGE works by sampling and aggregating features from a node’s immediate neighbors, thus reducing the computational load when handling large datasets. The core functionality of GraphSAGE is defined by its update rules, which specify how node representations are updated based on their neighborhoods:

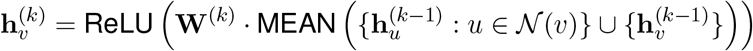

Here, 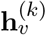 represents the feature vector of node *v* at layer *k*, 𝒩 (*v*) denotes the set of neighbors of *v*, and **W**^(*k*)^ are the trainable parameters at layer *k*. This allows scGRIP to efficiently aggregate and update gene and peak node features across multiple layers, enabling learning both the local regulatory modules and the gene/peak-specific information.

### Embedded Topic Model component

The Embedded Topic Model (ETM) [23] leverages an encoder-decoder framework, where the encoder maps high-dimensional data to a low-dimensional topic space, and the decoder reconstructs the original data from this topic representation, facilitating the discovery of underlying biological themes.

#### Encoder

The encoder is structured as a two-layer fully connected neural network, tasked with inferring topic proportions from normalized count vectors of multi-omics data for individual cells. It processes each cell’s multi-omics data through its layers to generate parameters that define a logistic normal distribution for each omics type. These distributions are assumed to capture the latent representation of the respective omics data, where each dimension corresponds to a potential topic or underlying biological factor. The primary goal of the encoder is to effectively synthesize these independent logistic normal distributions into a cohesive joint distribution that represents the comprehensive latent profile of the multi-omics data. This synthesis allows the model to capture the complex interdependencies and unique contributions of each omics layer.

Each modality has a dedicated encoder that produces the mean and variance of a *K*-dimensional latent Gaussian distribution. For modality *m* ∈ {gene, peak}:

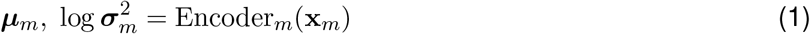

specifically, a multi-layer perceptron (MLP) architecture is employed:

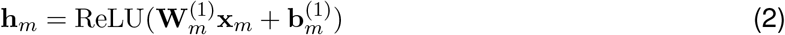

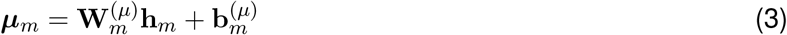

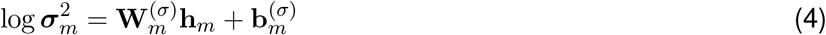

A reparameterization trick enables stochastic gradient descent through sampling:

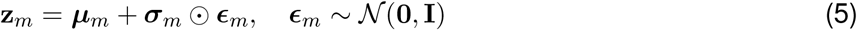

where

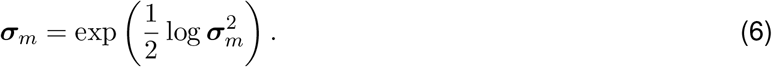

Latent topic mixture proportions are modeled as:

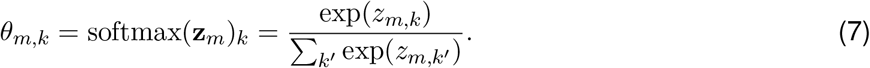

#### Decoder

The decoder in scGRIP leverages the embeddings produced by a GNN to reconstruct the input data from the topic embeddings and the node embeddings. The GNN produces cell-conditioned node embeddings **z**^(*c*)^ ∈ ℝ^*M* ×*L*^, which are shared between the decoder and downstream GRN inference.

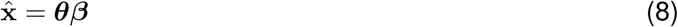

where

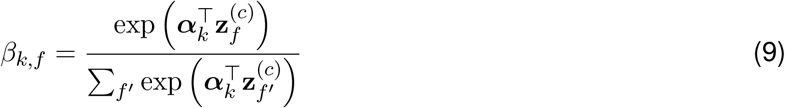

and 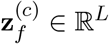 denotes the GNN-derived embedding of feature *f* in cell *c*. Here, *k* indexes topics, ***θ*** is the topic proportion vector, and ***β*** is the topic-feature matrix.

The optimization with respect to the encoder and decoder maximizes an evidence lower bound (ELBO) of the marginal log likelihood:

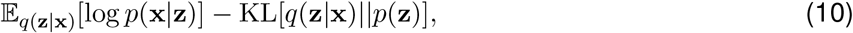

where the expectation is approximated by Monte Carlo sampling and the KL term has a closed-form expression:

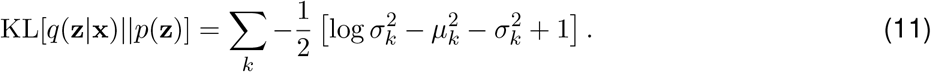

#### Model training and inference

The scGRIP model employs an end-to-end training approach where the encoder, decoder, and GNN are optimized simultaneously. The training is guided by a composite loss function that includes: average Negative Log-Likelihood (NLL), Kullback-Leibler (KL) divergence, and graph reconstruction loss. The overall loss function is formulated as:

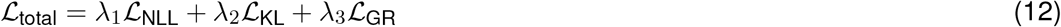

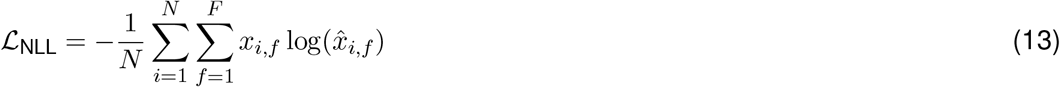

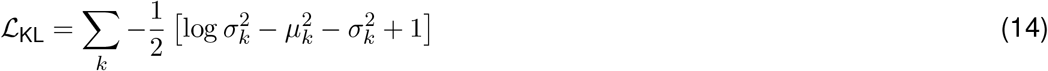

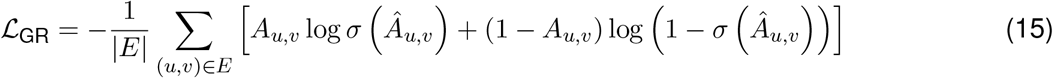

where *E* is the set of edges in the graph, **A** ∈ ℝ^*M* ×*M*^ is the adjacency matrix, and the reconstructed adjacency matrix is defined as:

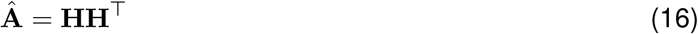

with **H** ∈ ℝ^*M* ×*L*^ denoting the concatenated node embeddings. *λ*_1_, *λ*_2_, and *λ*_3_ are dynamic weights used to balance the loss components during training [64].

### GRN inference

#### Predicting TF-RE links

The resulting node embeddings **z** ∈ ℝ^*M* ×*D*^, where *M* is the total number of features and *D* is the embedding dimension, produced by the GNN encode both global GRN topology and dynamic cell-specific context. The embeddings **z**^(*c*)^ are conditioned on cell-specific inputs, enabling dynamic, cell-resolved regulatory interaction modeling:

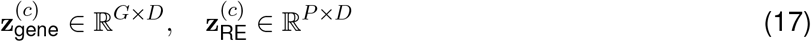

where *G* is the number of genes and *P* is the number of REs, with *G* + *P* = *M*.

To quantify cell-type-specific TF–RE regulatory interactions, we compute the average TF and RE embeddings across all cells belonging to a given cell type. The regulatory strength between transcription factor *t* and regulatory element *p* is then defined as the average of their dot product between embeddings:

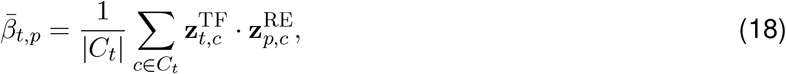

#### GraphSHAP

To compute GraphSHAP from GRN, we adapted GraphSVX [22]. In the original GraphSVX implementation, the explanation is local to a target node *v*. The method first extracts the *k*-hop neighborhood of *v* and defines the explanatory variables as the node features together with the neighboring nodes. It then samples coalitions *S* over this set. For each coalition, a perturbed input is constructed by masking excluded features and isolating excluded neighbors, and the original graph neural network is evaluated to obtain the prediction. Subsequently, GraphSVX fits a weighted linear surrogate model using Shapley kernel weights, where the estimated Shapley value associated with feature or neighbor quantifies its contribution to the prediction of node *v*. To compute edge attributions for RE–TG and TF–TG interactions, we adapt GraphSVX to operate on the regulatory graph derived from prior knowledge. Instead of explaining static node predictions, we condition the explanation on cell-specific inputs, allowing the attribution scores to capture dynamic, cell-resolved regulatory effects.

For a target gene *g*, we define its candidate regulator set as the union of transcription factors (TFs) and regulatory elements (REs) within its *k*-hop neighborhood:

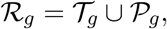

where 𝒯_*g*_ and 𝒫_*g*_ denote the TF and RE regulators of *g*, respectively.

Coalitions *S* ⊆ ℛ_*g*_ are sampled from this regulator set by masking subsets of regulators, implemented by removing their incident edges from the graph while keeping cell-specific features fixed. The model is then re-evaluated under each coalition to obtain perturbed predictions.

The prediction explained for gene *g* in cell *c* is given by

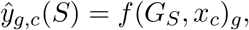

where *G*_*S*_ denotes the global regulatory graph after excluding regulators in ℛ_*g*_ \ *S*. The TF–TG and RE–TG Shapley values are then estimated by fitting a Kernel-SHAP style weighted linear surrogate over the coalition predictions:

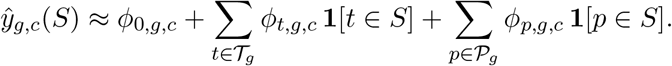

The surrogate is fitted using Shapley kernel weights

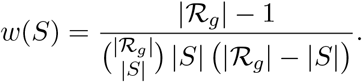

Here, *ϕ*_*t,g,c*_ represents the Shapley value corresponding to the TF–TG contribution of TF *t* to target gene *g* in cell *c*, while *ϕ*_*p,g,c*_ represents the RE–TG contribution of peak *p* on gene *g* in cell *c*. Positive values indicate that the regulator increases the predicted expression of *g* in cell *c*, whereas negative values indicate that it decreases the predicted expression.

#### Predicting RE-TG and TF-TG links

Let *t, p, g*, and *c* index TFs, REs (peaks), TGs, and cells, respectively. TF_*t,c*_ and RE_*p,c*_ denote cell-specific TF and RE activities, **z**^TF^, **z**^RE^, and **z**^TG^ denote learned node embeddings, and 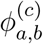 denotes the GraphSVX-derived Shapley attribution weight capturing the influence of node *a* on node *b* in cell *c*. Each coalition is encoded as a binary indicator vector describing which regulators are retained. Following GraphSVX, we fit a kernel-weighted linear surrogate model that maps coalition indicators to model predictions, where the regression coefficients approximate Shapley attribution values for each regulator.

The TF–RE association strength 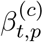 is defined as the embedding similarity between the TF and RE:

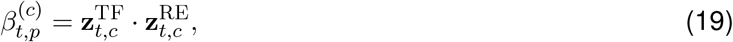

The RE–TG association strength 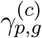 incorporates GraphSVX-derived SHAP attribution together with embedding similarity:

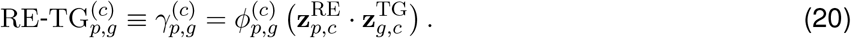

The resulting SHAP-adjusted per-cell scores define a cell-specific GRN representation for downstream significance analysis:

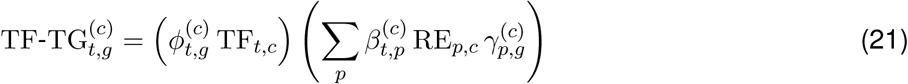

This procedure yields a weighted, cell-resolved GRN in which edges vary across cells according to regulatory activity rather than remaining fixed across the population or cell type.

#### Cell-type- and condition-specific GRN inference

Cells belonging to the target cell type are defined as the target set, while all remaining cells serve as the background. For each regulatory pair, we compare the distribution of GraphSVX-derived per-cell regulatory scores (Eq. 21) between the two groups using the Wilcoxon rank-sum test. The resulting *p*-values are adjusted for multiple testing using the Benjamini–Hochberg procedure, and regulatory pairs with adjusted *p*-value *<* 0.05 are retained as significant interactions, forming the cell-type-specific GRN.

To characterize condition-dependent regulation, we perform a second statistical test within each cell type. Specifically, for a given cell type, cells from the focal condition are defined as the target set, while cells from all other conditions of the same cell type serve as the background. For each regulatory pair, we again compare the distribution of GraphSVX-derived per-cell regulatory scores between the two groups using the Wilcoxon rank-sum test, followed by Benjamini–Hochberg correction. Significant regulatory pairs identified under this contrast capture condition-specific regulatory changes within each cell type.

### Single-cell multi-omic benchmark data

1. Peripheral Blood Mononuclear Cells (PBMC) from 10X Genomics, consisting of 9,631 cells, 29,095 genes, and 112,975 peaks across 19 cell types.
2. Multiome bone marrow mononuclear cells (BMMC) dataset from the 2021 NeurIPS challenge [65], consisting of 69,249 cells, 13,403 genes and 110,359 peaks with 22 cell types from 10 donors across 4 sites
3. Human Cortex dataset from BRAIN Initiative [66, 67], consisting of 45,549 cells, 30,033 genes, 262,997 peaks with 13 cell types.
4. Multiome AD vs Control DLPFC [15], consisting 105,332 nuclei obtained from 15 individuals (7 Alzheimer’s disease and 8 controls) with 189,925 ATAC peaks in total.

All datasets were processed into the format of samples-by-features matrices. Initially, the read count for each gene and peak were first normalized per cell by total counts of 1 × 10^4^ within the same omic using the `scanpy.pp.normalize_total` function in Scanpy. Next, `log1p` transformation was applied. The `scanpy.pp.highly_variable_genes` function was used to select highly variable genes or peaks for downstream analyses; clustering and imputation tasks were allowed to operate with or without HVG selection, whereas all other tasks required HVG-filtered features. Peaks were selected following the GRN graph construction steps based on TSS and CisTarget.

### Method evaluation

#### TF-RE inference evaluation

To evaluate our model’s ability to predict cell-type level TF-RE interactions, we used ChIP-seq atlas data [25] as ground truth. For each TF *t*, we constructed a balanced evaluation set defined as whether each peak *p* is bound by TF *t* according to the ChIP-seq experiment. For the negative samples, we used the equal number of randomly selected negative regions. Using these balanced positive and negative sets, we calculate the Area Under the Precision-Recall Curve (AUPRC) as the evaluation metric to assess the overall balance between precision and recall.

#### Cell-type or condition classification by the inferred GRN

To evaluate whether the inferred cell-specific GRNs capture discriminative regulatory signatures, we formulate a cell-type classification task. For each cell type, we construct cell-level feature representations using the regulatory scores (Eq. 21) restricted to the significant regulatory pairs identified for that cell type. Each cell is labeled by its cell-type annotation, or by supertype if subtype merging is enabled, and the task is to predict whether a cell belongs to the focal cell type versus all other cell types.

To focus on evaluating the inferred GRN rather than the classifier itself, we employ a *k*-nearest neighbors (KNN) classifier, which operates directly on the constructed feature space without requiring parametric training. Hyperparameters, including the number of neighbors, distance metric, feature scaling, and weighting scheme, are selected via grid search on the validation split. The final model is evaluated on a held-out test split, and performance is summarized using AUROC (**Table** S6).

We perform an analogous evaluation for condition classification. For each cell type, we construct feature representations using the corresponding significant regulatory pairs and predict condition labels (e.g., AD versus control). Since cell-level condition annotations are not available, labels are assigned based on subject-level condition information (**Table** S7).

#### Reproducibility analysis of the AD differential GRN

To confirm that the significant pairs identified by scGRIP are reproducible across studies, we validated the method on an independent AD dataset [46]. For each overlapping cell type, we repeated the entire procedure of scoring, statistical testing, and multiple testing correction to identify significant TF-RE element pairs. Reproducibility was quantified using hypergeometric testing, which measures the overlap between significant pairs derived from the primary and validation datasets. Because the two datasets have distinct definitions of chromatin accessibility peaks, we mapped peaks between datasets using bedtools, considering peaks matched if they overlapped by at least one base pair. The same procedure was applied to compare the reproducibility of differentially expressed genes and differentially accessible regions.

#### Cell clustering

Using the cell topic distribution **θ** from the scGRIP encoder, we applied the Louvain algorithm [68] to cluster cells. To evaluate performance on the cell type clustering task, we employed four evaluation metrics to assess performance:

1. Adjusted Rand Index (ARI) [69]: quantifies the similarity between two clusters while correcting for the chance that pairs of objects might be randomly assigned to the same clusters.
2. Normalized Mutual Information (NMI) [70]: measures the amount of information shared between two clusters, normalized by the average entropy of the clusters.
3. kBET [71]: tests whether batch labels are distributed differently across cells using Pearson’s chisquare test.
4. GC: evaluates whether cells of the same type from different batches are close in the embedding space by constructing a k-nearest neighbor graph.

#### Cross-modality imputation

To evaluate the cross-modality imputation task, we compute the Pearson correlation coefficients (PCC) and Spearman correlation between the predicted and observed signals. Experiments were conducted on both highly variable genes and coding genes, whereas for GRN inference, evaluation was performed using highly variable genes only. Hyperparameters can be found in supplementary table S4 and S5

### Baseline methods

As the first baseline method for GRN inference, we evaluated LINGER version *1*.*105*. The parameters passed to LingerGRN were *genome=hg38, method=LINGER, activef=ReLu, singlepseudobulk=0*, and cell specific scores are outputed via:

#### TF–RE

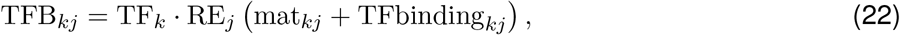

where mat_*kj*_ denotes the TF–RE binding strength, TFbinding_*kj*_ denotes motif binding affinity and RE_*j*_ and TG_*i*_ are cell specific expressions.

#### RE–TG

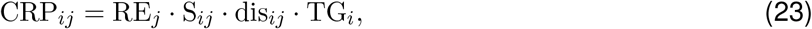

where S_*ij*_ represents the SHAP-derived cis-regulatory strength, dis_*ij*_ is the genomic distance weighting, and RE_*j*_ and TG_*i*_ are cell specific expressions.

#### TF–TG

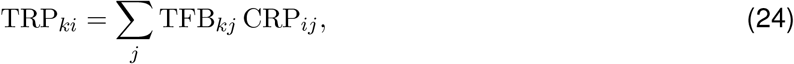

As the second baseline method, we evaluated GLUE (scglue) version *0*.*4*.*1*. Parameters for GLUE were kept at defaults unless otherwise noted (e.g., *lam_data=1*.*0, lam_kl=1*.*0, lam_graph=0*.*02, lam_align=0*.*05, lam_sup=0*.*02, dsc_steps=1, lr=2e-3*). Training used the default graph minibatching scheme with *graph_batch _size=32*. The guidance graph was created from our prior GRN matrix, with all edges assigned *weight=1*.*0* and *sign=1*. GRN inference was performed using `scglue.genomics.regulatory_inference` to obtain feature-feature scores. Cell-type–specific GRNs were derived by weighting the TF–TG, RE–TG, TF–RE edges by cell-type mean expression.

## Declarations

### Availability of data and materials

All datasets used in this study are publicly available and were obtained from established data repositories.The single-cell multimodal datasets used in this study are as follows: the PBMC dataset is available at 10X Genomics https://www.10xgenomics.com/datasets/10-k-peripheral-blood-mononuclear-cells-pbm-cs-from-a-healthy-donor-1-standard-1-1-0, the BMMC dataset and the Cerebral Cortex dataset can be found on Gene Expression Omnibus under series numbers GSE194122 and GSE204684, respectively. The AD dataset can be found under GSE214637. For the gene regulatory network databases, cisTarget is available at https://resources.aertslab.org/cistarget/, and the ChIP-seq Atlas is accessible at https://chip-atlas.org/. All original code has been deposited at https://github.com/li-lab-mcgill/scGRIP.

### Authors’ contributions

Y.L. conceptualized the study and designed the experiment. W.D. and M.Z. implemented the scGRIP and performed the experiments under the supervision of Y.L. All authors read and approved the final manuscript.

## Acknowledgment

We thank members of the Li lab for their feedback and comments on earlier iterations of this work.

