## Supplemental tables and figures for "scGRIP: a graph-based explainable AI framework for single-cell multi-omics Gene Regulatory Inference with Prior Knowledge"

### Supplementary Tables

| Cell Type | TF | scGRIP | LINGER | GLUE | PCC |
|---|---|---|---|---|---|
| CD4 T cell | FOS | 0.610 | 0.600 | 0.600 | 0.595 |
| CD4 T cell | IRF4 | 0.635 | 0.595 | 0.590 | 0.595 |
| CD4 T cell | BCL6 | 0.650 | 0.520 | 0.615 | 0.605 |
| CD4 T cell | FOXP3 | 0.615 | 0.625 | 0.605 | 0.605 |
| CD4 T cell | FOSL1 | 0.610 | 0.620 | 0.640 | 0.622 |
| CD4 T cell | BATF | 0.580 | 0.590 | 0.590 | 0.602 |
| CD4 T cell | GATA3 | 0.560 | 0.580 | 0.565 | 0.575 |
| CD4 T cell | RUNX1 | 0.565 | 0.565 | 0.590 | 0.565 |
| CD4 T cell | ETS1 | 0.620 | 0.620 | 0.630 | 0.605 |
| CD4 T cell | Average | **0.630** | 0.620 | 0.625 | 0.615 |
| B cell | BACH2 | 0.590 | 0.585 | 0.600 | 0.560 |
| B cell | BCL6 | 0.650 | 0.580 | 0.620 | 0.590 |
| B cell | CTCF | 0.620 | 0.625 | 0.625 | 0.596 |
| B cell | EBF1 | 0.580 | 0.560 | 0.600 | 0.575 |
| B cell | IRF4 | 0.630 | 0.600 | 0.625 | 0.595 |
| B cell | MEF2B | 0.585 | 0.525 | 0.555 | 0.565 |
| B cell | Average | **0.610** | 0.585 | 0.605 | 0.585 |
| Memory B cell | IRF4 | **0.620** | 0.615 | 0.605 | 0.565 |

Table S1: AUPRC for cell type-specifc TF-RE prediction on the PBMC dataset. For each cell type, the values are averaged across TFs, with the best performance highlighted in bold.

| Dataset | Method | ARI | NMI | kBET | GC |
|---|---|---|---|---|---|
| PBMC | scGRIP | **0.900 (0.024)** | **0.885 (0.026)** | - | - |
| | scGRIP _allCodingGene | 0.860 (0.022) | 0.861 (0.012) | - | - |
| | scGRIP _noGraph | 0.810 (0.053) | 0.823 (0.043) | - | - |
| | MultiVI | 0.801 (0.029) | 0.811 (0.024) | - | - |
| | Seurat | 0.725 (0.051) | 0.791 (0.013) | - | - |
| | MoETM | 0.828 (0.019) | 0.831 (0.016) | - | - |
| BMMC | scGRIP | **0.733 (0.021)** | **0.797 (0.020)** | **0.251 (0.023)** | **0.942 (0.005)** |
| | scGRIP _allCodingGene | 0.623 (0.035) | 0.780 (0.023) | 0.181 (0.007) | 0.877 (0.006) |
| | scGRIP _noGraph | 0.242 (0.011) | 0.748 (0.030) | 0.242 (0.011) | 0.925 (0.004) |
| | MultiVI | 0.714 (0.021) | 0.756 (0.024) | 0.211 (0.009) | 0.941 (0.006) |
| | Seurat | 0.605 (0.021) | 0.732 (0.011) | 0.136 (0.004) | 0.847 (0.006) |
| | MoETM | 0.709 (0.013) | 0.783 (0.007) | 0.241 (0.018) | 0.926 (0.003) |
| Cerebral Cortex | scGRIP | **0.689 (0.029)** | **0.783 (0.025)** | 0.221 (0.015) | 0.934 (0.006) |
| | scGRIP _allCodingGene | 0.650 (0.042) | 0.729 (0.023) | 0.187 (0.011) | 0.909 (0.004) |
| | scGRIP _noGraph | 0.642 (0.038) | 0.689 (0.023) | 0.221 (0.021) | 0.912 (0.005) |
| | MultiVI | 0.608 (0.034) | 0.701 (0.011) | 0.188 (0.004) | 0.944 (0.005) |
| | Seurat | 0.549 (0.030) | 0.728 (0.014) | 0.171 (0.006) | 0.909 (0.008) |
| | MoETM | 0.644 (0.011) | 0.779 (0.007) | **0.287 (0.025)** | **0.937 (0.004)** |

Table S2: **Evaluation of cell clustering.** Under each dataset, the best score per evaluation metric is shown in bold, and the second-best score is shown in blue. Values represent mean (standard deviation).

| Dataset | Method | ATAC2RNA | | RNA2ATAC | |
|---|---|---|---|---|---|
| | | Pearson Corr | Spearman Corr | Pearson Corr | Spearman Corr |
| PBMC | scGRIP | **0.87 (0.05)** | 0.36 (0.02) | 0.57 (0.04) | 0.34 (0.02) |
| | scGRIP _allCodingGene | 0.64 (0.03) | 0.31 (0.02) | 0.53 (0.03) | **0.36 (0.03)** |
| | scGRIP _impute | 0.82 (0.04) | 0.37 (0.01) | **0.58 (0.02)** | 0.35 (0.02) |
| | scGRIP _allCodingGene_impute | 0.67 (0.02) | 0.35 (0.02) | 0.57 (0.02) | 0.35 (0.02) |
| | Babel | 0.63 (0.05) | 0.34 (0.01) | 0.53 (0.02) | 0.32 (0.02) |
| | MultiVI | 0.76 (0.02) | 0.39 (0.01) | 0.57 (0.01) | 0.35 (0.01) |
| | MoETM | 0.78 (0.03) | **0.45 (0.02)** | 0.49 (0.03) | 0.33 (0.04) |
| BMMC | scGRIP | 0.66 (0.04) | **0.41 (0.03)** | 0.57 (0.03) | 0.36 (0.02) |
| | scGRIP _allCodingGene | 0.61 (0.04) | 0.34 (0.04) | 0.55 (0.04) | 0.35 (0.01) |
| | scGRIP _impute | **0.69 (0.04)** | 0.41 (0.02) | 0.58 (0.03) | **0.37 (0.02)** |
| | scGRIP _allCodingGene_impute | 0.59 (0.01) | 0.36 (0.03) | 0.58 (0.02) | 0.36 (0.02) |
| | Babel | 0.61 (0.02) | 0.38 (0.03) | 0.52 (0.02) | 0.32 (0.03) |
| | MultiVI | 0.63 (0.02) | 0.4 (0.01) | 0.58 (0.02) | 0.34 (0.01) |
| | MoETM | 0.63 (0.02) | 0.39 (0.01) | **0.59 (0.02)** | 0.33 (0.04) |
| Cerebral Cortex | scGRIP | 0.64 (0.03) | 0.37 (0.02) | 0.56 (0.02) | **0.37 (0.03)** |
| | scGRIP _allCodingGene | 0.67 (0.05) | 0.35 (0.02) | 0.56 (0.05) | 0.35 (0.04) |
| | scGRIP _impute | 0.67 (0.04) | **0.42 (0.02)** | 0.55 (0.02) | 0.33 (0.01) |
| | scGRIP _allCodingGene_impute | **0.68 (0.03)** | 0.36 (0.01) | 0.55 (0.02) | 0.36 (0.02) |
| | Babel | 0.66 (0.03) | 0.37 (0.02) | 0.53 (0.01) | 0.31 (0.02) |
| | MultiVI | 0.64 (0.01) | 0.42 (0.02) | 0.56 (0.02) | 0.36 (0.01) |
| | MoETM | 0.63 (0.02) | 0.4 (0.01) | **0.59 (0.02)** | 0.33 (0.01) |

Table S3: **Cross-modality imputation evaluation.** We imputed gene expression values from chromatin accessibility values (ATAC2RNA) and vice versa (RNA2ATAC). Under each dataset, the best score per evaluation metric in each direction is in bold, and the second best score is in blue. The values represent the mean (standard deviation).

| **Learning Rate** | | **Embedding Size** | | **Topic Number** | |
|---|---|---|---|---|---|
| **Value** | **Train ARI** | **Value** | **Train ARI** | **Value** | **Train ARI** |
| 1e-3 | [0.745] | 128 | [0.812] | 20 | [0.849] |
| 1e-4 | [0.883] | 256 | [0.843] | 40 | [0.862] |
| 1e-5 | [0.711] | 512 | [0.883] | 60 | [0.858] |
| | | | | 80 | [0.854] |
| | | | | 100 | [0.877] |
| **TSS Threshold** | | **HVG** | | **Latent Size** | |
| **Value** | **Train ARI** | **Value** | **Train ARI** | **Value** | **Train ARI** |
| 150e3 | [0.876] | 2000 | [0.837] | 10 | [0.677] |
| 250e3 | [0.872] | 3000 | [0.848] | 50 | [0.723] |
| 1e6 | [0.870] | 4000 | [0.853] | 100 | [0.831] |
| | | 5000 | [0.849] | | |

Table S4: **scGRIP Hyperparameters selection based on ARI Performance on PBMC dataset**. We conducted an extensive grid search over key hyperparameters, reporting the resulting Adjusted Rand Index (ARI) for each configuration. The learning rate of 1e-4, embedding size of 512, and topic number of 100 yielded the best overall clustering performance. Additionally, we evaluated the impact of technical parameters such as TSS (Transcription Start Site) threshold, number of highly variable genes (HVG), and latent dimensions.

| **Hyperparameter** | **Values** | **Selected Value** |
|---|---|---|
| Learning Rate | {1e-2, 1e-3, 1e-4 } | 1e-2 |
| Embedding Size | {128, 256, 512} | 512 |
| p | {0.25, 0.5, 0.75, 1.0} | 0.5 |
| q | {1.0, 2.0, 3.0, 4.0} | 2.0 |
| Walk Length | {10, 20, 40, 80} | 20 |
| Number of Walks | {10, 20, 30, 50} | 30 |
| Window Size | {5, 10, 15, 20} | 10 |
| Iterations | {5, 10, 20, 50} | 10 |

Table S5: **Node2Vec Hyperparameters and Training Settings**. The exploration-exploitation parameters p and q were tuned to balance local and global network structures, while walk length, number of walks, and window size were optimized to capture meaningful connectivity patterns in the biological networks. The selected configuration provided the best trade-off between computational efficiency and representation quality.

Table S6: Cell-type prediction AUROC across regulatory relationships.

| Regulatory Type | Cell Type | scGRIP | scGRIP (SHAP Ablation) | scGRIP (All Coding Gene) | LINGER |
|---|---|---|---|---|---|
| TF–RE | Astrocytes | 0.873 ± 0.042 | – | 0.874 ± 0.024 | 0.889 ± 0.020 |
| | Inhibitory | 0.872 ± 0.028 | – | 0.871 ± 0.022 | 0.883 ± 0.025 |
| | Excitatory | 0.885 ± 0.035 | – | 0.868 ± 0.020 | 0.904 ± 0.018 |
| | Microglia | 0.906 ± 0.030 | – | 0.880 ± 0.026 | 0.901 ± 0.033 |
| | OPC | 0.925 ± 0.025 | – | 0.882 ± 0.021 | 0.895 ± 0.022 |
| | Oligodendrocytes | 0.886 ± 0.018 | – | 0.873 ± 0.019 | 0.887 ± 0.023 |
| RE–TG | Astrocytes | 0.937 ± 0.034 | 0.859 ± 0.048 | 0.922 ± 0.028 | 0.929 ± 0.037 |
| | Inhibitory | 0.940 ± 0.026 | 0.862 ± 0.030 | 0.911 ± 0.021 | 0.933 ± 0.028 |
| | Excitatory | 0.950 ± 0.021 | 0.866 ± 0.035 | 0.914 ± 0.018 | 0.912 ± 0.037 |
| | Microglia | 0.953 ± 0.026 | 0.860 ± 0.038 | 0.923 ± 0.024 | 0.929 ± 0.047 |
| | OPC | 0.921 ± 0.034 | 0.831 ± 0.037 | 0.904 ± 0.019 | 0.918 ± 0.047 |
| | Oligodendrocytes | 0.926 ± 0.021 | 0.867 ± 0.039 | 0.908 ± 0.018 | 0.896 ± 0.039 |
| TF–TG | Astrocytes | 0.953 ± 0.035 | 0.898 ± 0.044 | 0.905 ± 0.030 | 0.940 ± 0.033 |
| | Inhibitory | 0.958 ± 0.044 | 0.911 ± 0.033 | 0.926 ± 0.024 | 0.931 ± 0.044 |
| | Excitatory | 0.933 ± 0.036 | 0.901 ± 0.044 | 0.901 ± 0.025 | 0.938 ± 0.035 |
| | Microglia | 0.951 ± 0.048 | 0.888 ± 0.036 | 0.894 ± 0.026 | 0.937 ± 0.044 |
| | OPC | 0.929 ± 0.036 | 0.903 ± 0.036 | 0.905 ± 0.019 | 0.928 ± 0.055 |
| | Oligodendrocytes | 0.941 ± 0.033 | 0.900 ± 0.045 | 0.908 ± 0.021 | 0.921 ± 0.047 |

Table S7: Disease prediction AUROC across regulatory relationships.

| Regulatory Type | Cell Type | scGRIP | scGRIP (SHAP Ablation) | scGRIP (All Codign Gene) | LINGER |
|---|---|---|---|---|---|
| TF–RE | Astrocytes | 0.711 ± 0.041 | – | 0.692 ± 0.028 | 0.728 ± 0.129 |
| | Inhibitory Neuron | 0.705 ± 0.047 | – | 0.688 ± 0.030 | 0.702 ± 0.137 |
| | Excitatory Neuron | 0.724 ± 0.032 | – | 0.698 ± 0.024 | 0.722 ± 0.152 |
| | Microglia | 0.720 ± 0.043 | – | 0.696 ± 0.031 | 0.719 ± 0.097 |
| | OPC | 0.694 ± 0.036 | – | 0.684 ± 0.022 | 0.714 ± 0.141 |
| | Oligodendrocytes | 0.710 ± 0.023 | – | 0.691 ± 0.021 | 0.706 ± 0.113 |
| RE–TG | Astrocytes | 0.754 ± 0.047 | 0.728 ± 0.049 | 0.720 ± 0.033 | 0.755 ± 0.036 |
| | Inhibitory | 0.757 ± 0.045 | 0.730 ± 0.031 | 0.721 ± 0.029 | 0.758 ± 0.027 |
| | Excitatory | 0.765 ± 0.022 | 0.733 ± 0.034 | 0.722 ± 0.024 | 0.741 ± 0.038 |
| | Microglia | 0.767 ± 0.044 | 0.728 ± 0.039 | 0.721 ± 0.031 | 0.755 ± 0.046 |
| | OPC | 0.741 ± 0.033 | 0.714 ± 0.038 | 0.712 ± 0.023 | 0.746 ± 0.048 |
| | Oligoden | 0.745 ± 0.023 | 0.725 ± 0.037 | 0.716 ± 0.020 | 0.728 ± 0.041 |
| TF–TG | Astrocytes | 0.763 ± 0.042 | 0.730 ± 0.051 | 0.722 ± 0.034 | 0.759 ± 0.053 |
| | Inhibitory Neuron | 0.767 ± 0.038 | 0.734 ± 0.041 | 0.721 ± 0.028 | 0.751 ± 0.039 |
| | Excitatory Neuron | 0.766 ± 0.052 | 0.733 ± 0.048 | 0.730 ± 0.032 | 0.725 ± 0.071 |
| | Microglia | 0.748 ± 0.041 | 0.725 ± 0.039 | 0.738 ± 0.027 | 0.763 ± 0.037 |
| | OPC | 0.763 ± 0.038 | 0.731 ± 0.032 | 0.719 ± 0.022 | 0.739 ± 0.087 |
| | Oligodendrocytes | 0.762 ± 0.027 | 0.729 ± 0.038 | 0.728 ± 0.021 | 0.714 ± 0.103 |

### Supplementary Figures

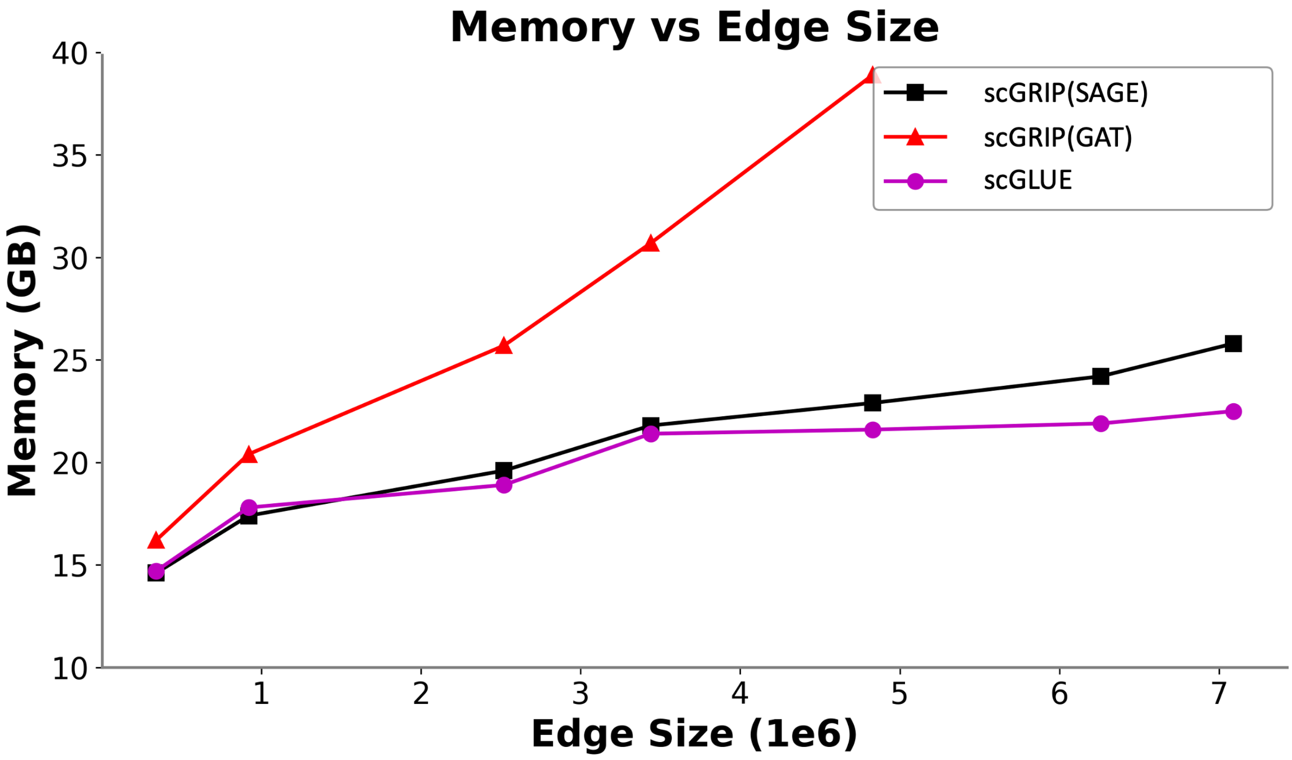


Figure S1: **Memory consumption as the input graph size increases**. This figure compares the computational efficiency of scGRIP against other graph-based approaches. Our GraphSAGE-based implementation demonstrates favorable scaling characteristics, showing comparable memory efficiency.

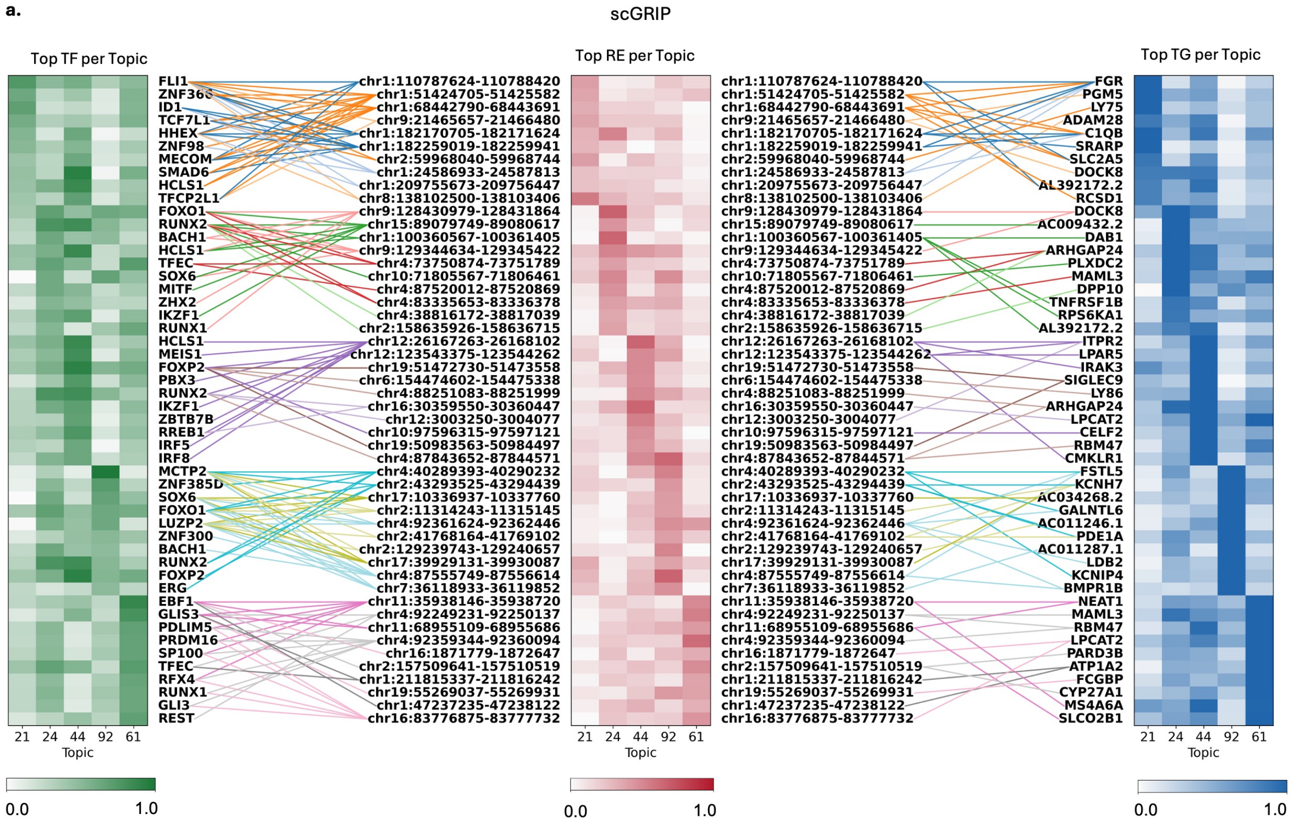


Figure S2: **Topic analysis reveals enriched regulatory modules in the AD dataset**. **a.** Feature analysis for the selected topics. Heatmaps display the top 10 transcription factors (TFs), regulatory elements (REs), and target genes (TGs) ranked by topic weight. TF–RE–TG regulatory triplets identified within each topic are highlighted with colored connections, illustrating topic-specific predicted regulatory modules captured by scGRIP. Bold labels denote TFs, REs, and TGs that participate in the identified regulatory triplets.

**a.**

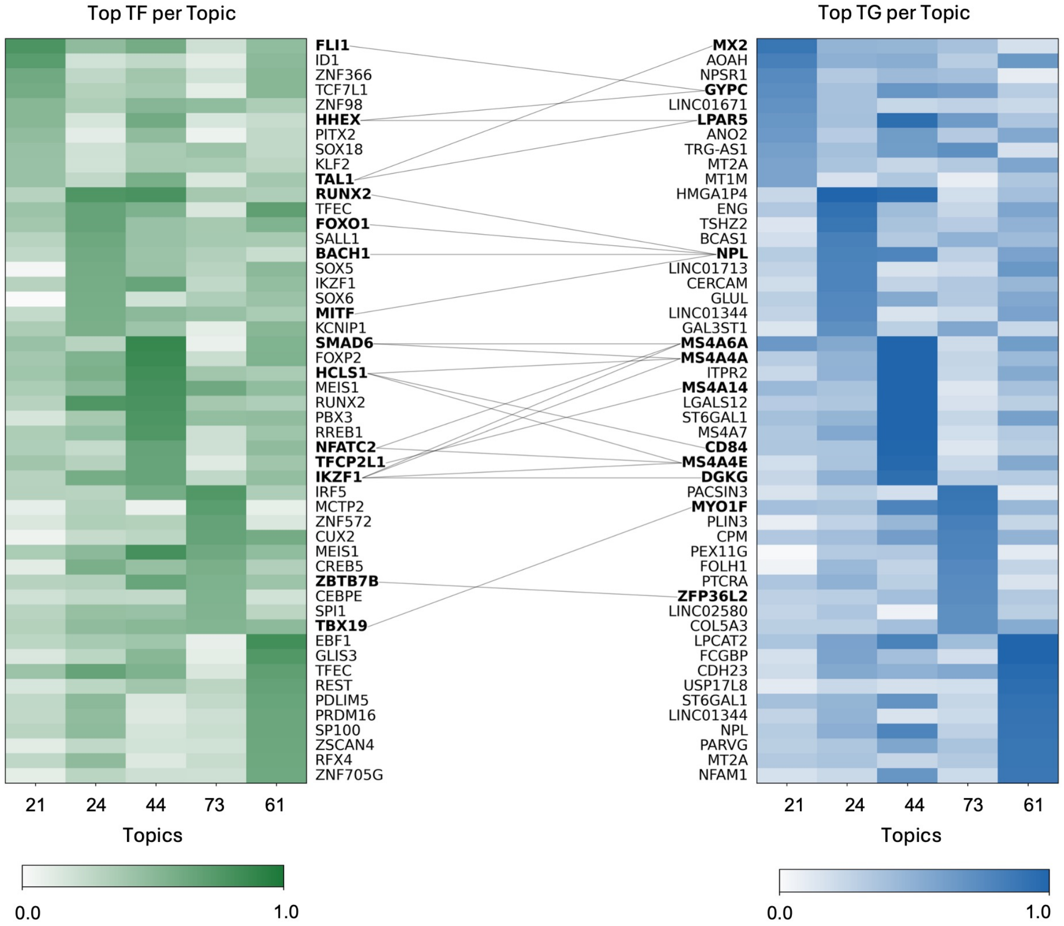


Figure S3: **Topic analysis reveals predited TF-TG connections in the AD dataset**. **a.** Feature analysis for the selected topics. Heatmaps display the top 10 transcription factors (TFs), and target genes (TGs) ranked by topic weight. TF–TG regulatory pair connections within each topic are calculated from the average TF–TG scores of the top 10% weighted cells in each topic. Bold labels denote TF and TGs gene names.

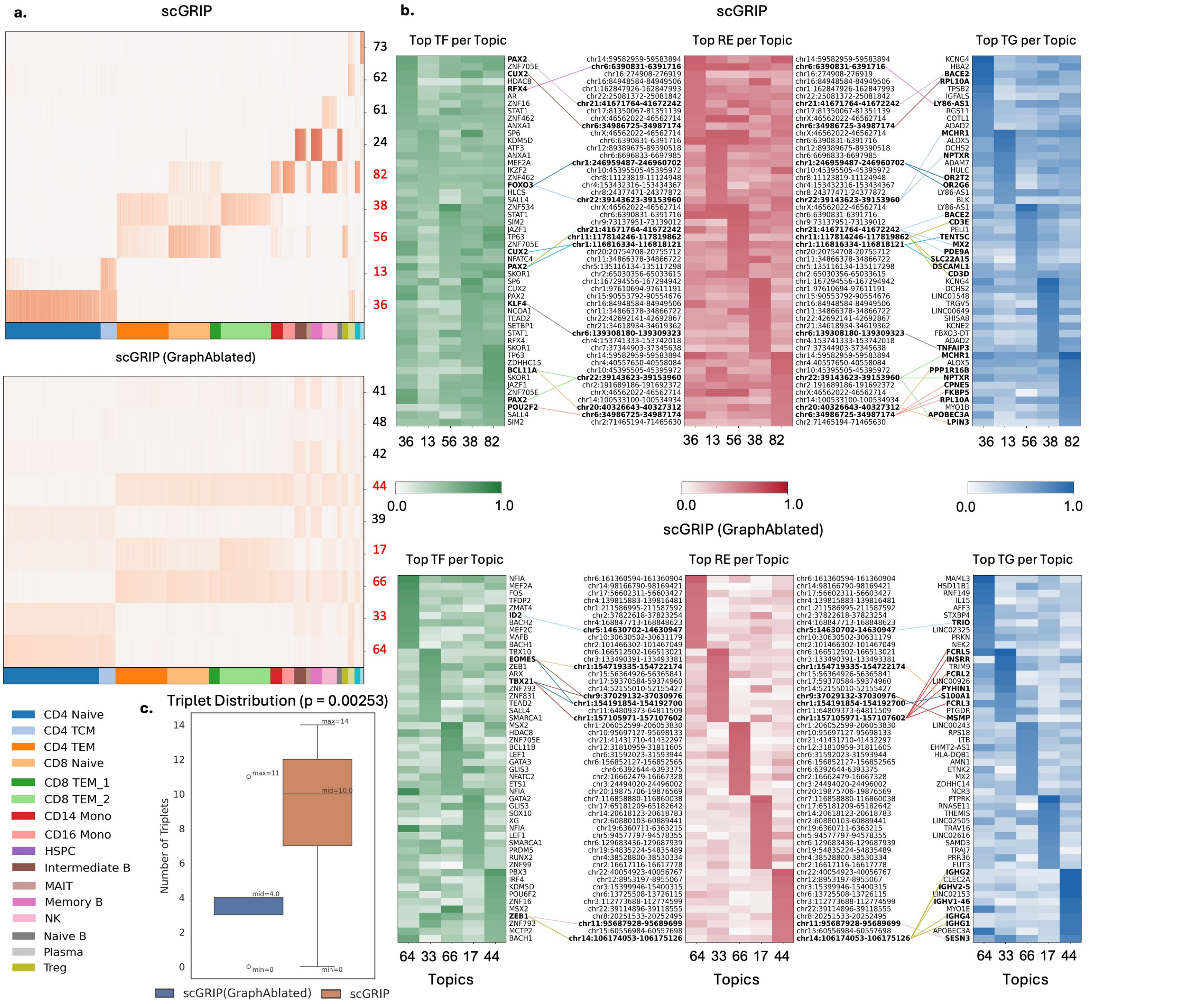


Figure S4: **Topic analysis reveals enriched regulatory modules in the PBMC dataset**. **a.** Heatmaps showing the intensity of the top topics from the PBMC dataset using scGRIP (top) and scGRIP (Graph-ablated) (bottom) across cell types. Topics highlighted in red correspond to topics in panel b. **b.** Feature analysis for the selected topics. Heatmaps display the top 10 transcription factors (TFs), regulatory elements (REs), and target genes (TGs) ranked by topic weight. TF–RE–TG regulatory triplets identified within each topic are highlighted with colored connections, illustrating topic-specific regulatory modules captured by scGRIP. Bold labels denote TFs, REs, and TGs that participate in the identified regulatory triplets. **c.** Comparison of the number of regulatory triplets identified among the top features per topic between scGRIP and the graph-ablated variant. scGRIP identifies a greater number of regulatory triplets, indicating improved recovery of coherent regulatory modules when incorporating graph-based regulatory priors.
